# Measuring host immune response status by simultaneous quantitative measurement of activity of signal transduction pathways that coordinate functional activity of immune cells from innate and adaptive immune system

**DOI:** 10.1101/2021.10.06.463309

**Authors:** Wilbert Bouwman, Wim Verhaegh, Arie van Doorn, Anja van de Stolpe

## Abstract

For many diseases, including cancer, viral infections such as COVID-19, bacterial infections, and auto-immune diseases, the immune response is a major determinant of progression, response to therapy, and clinical outcome. Innate and adaptive immune response are controlled by coordinated activity of multiple immune cell types. The functional activity state of immune cells is determined by cellular signal transduction pathways (STPs). A novel mRNA-based signaling pathway assay platform has been developed to quantitatively measure relevant STP activities in all types of immune cells and mixed immune cell samples for experimental and diagnostic purposes. We generated a STP activity profile, termed Immune-Pathway Activity Profile (I-PAP), for a variety of immune cell types in resting and activated state, and provide a first example for use in patient samples.

**Methods:** The technology to measure STP activity has been described for androgen and estrogen receptor, PI3K, MAPK, TGFβ, Notch, NFκB, JAK-STAT1/2, and JAK-STAT3 pathways. STP activity was measured on Affymetrix expression microarray data from preclinical studies containing public data from different types of immune cells, resting/naïve or immune-activated *in vitro*, to establish I-PAPs. Subsequently data from a clinical study on rheumatoid arthritis were analyzed.

**Results:** I-PAPs of naïve/resting and immune-activated CD4+ and CD8+ T cells, T helper cells, B cells, NK cells, monocytes, macrophages, and dendritic cells were established and in agreement with known experimental immunobiology. In whole blood samples of rheumatoid arthritis patients TGFβ pathway activity was increased; JAK-STAT3 pathway activity was selectively increased in female patients. In naïve CD4+ Tregs TGFβ pathway activity was increased, while in memory T effector cells JAK-STAT3 pathway activity tended to increase, suggesting that these immune cell types contributed to whole blood analysis results.

**Conclusion:** STP assay technology (currently being converted to qPCR-based assays) makes it possible to directly measure functional activity of cells of the innate and adaptive immune response enabling quantitative assessment of the immune response of an individual patient. Envisioned utility lies in (1) prediction and monitoring of response to immunomodulatory treatments for a variety of immune-mediated diseases, including RA; (2) uncovering novel treatment targets; (3) improvement and standardization of *in vitro* immunology research and drug development.

## Introduction

The immune response is determined by the coordinated activity of a number of immune cell types belonging to the innate or adaptive immune system (Davis, n.d.) (Figure 1A). Immune cells communicate through a variety of ligands that bind specifically to receptors on the cell membrane or inside the cell (Figure 1B). Upon ligand binding, receptors specifically activate signal transduction pathways (STPs), which through intracellular crosstalk may activate other STPs, resulting in highly controlled changes in cell function (Cantley et al., 2014). Every immune cell type has its own set of membrane receptors and signal transduction pathways (STPs) determining to which signals the cell will respond. Growth hormones, cytokines, interferons, estrogens and androgens, are signals that act in concert to control activity of STPs such as PI3K- FOXO, MAPK, JAK-STAT1/2 and JAK-STAT 3, NFκB, TGFβ, Notch, estrogen (ER) and androgen (AR) receptor pathways, to regulate functions such as immune cell maturation and activation (Han et al., 2012),(Platanias, 2005),(Oh and Ghosh, 2013),(Li et al., 2007),(Goodman et al., 2011),(Perchet et al., 2018),(Deftos and Bevan, 2000),(Liu et al., 2007),(Lai et al., 2012),(Kovats, 2015). Ligand activated STP activity can be regulated in a ligand-dependent manner or through intracellular crosstalk between STPs.

**Figure 1.**
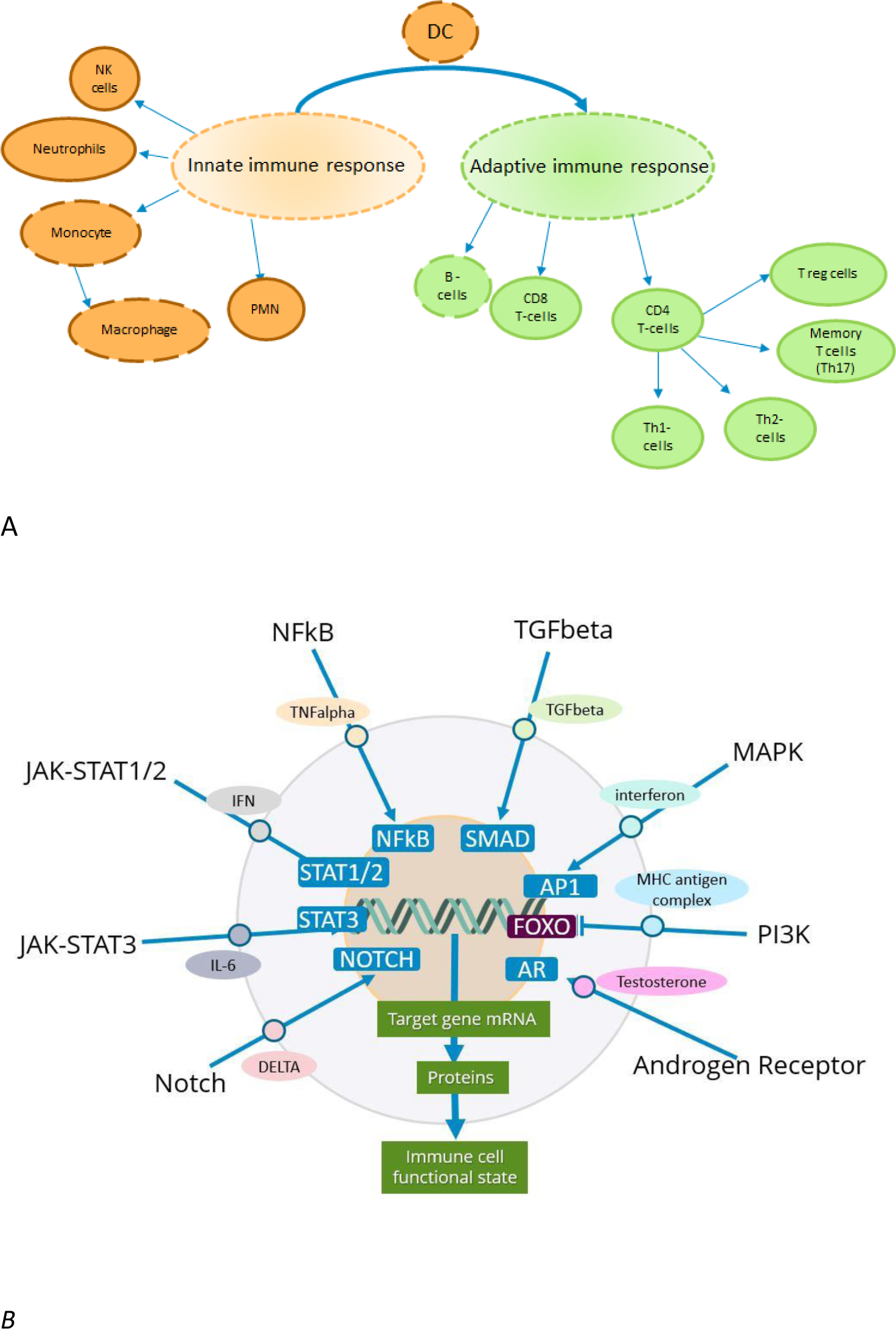
A. Summary of the immune cell types that play a role in the innate and adaptive immune response and are characterized in this study. Dashed line around cell type indicates antigen-presenting cell type. B. Simplified representation of an immune cell with relevant signal transduction pathways, illustrative ligand (oval), receptor (circle), and transcription factor (blue rectangle). Every immune cell type may have its own set of activatable signal transduction pathways.

Using a previously reported novel assay technology, we measured activity of immunologically relevant STPs in different types of immune cells, in naïve or resting and immune-activated state. Based on the results, we determined an Immune-signal transduction Pathway Activity Profile (I- PAP) for each cell type, in the two different functional states. To explore clinical value of the assay technology, STP activity was measured in blood samples of patients with rheumatoid arthritis (RA).

## Methods

### Measurement of activity of signal transduction pathways on Affymetrix microarray data from cell samples

Development and validation of Affymetrix-based assays for measuring activity of the estrogen (ER) and androgen receptor (AR), PI3K-FOXO, MAPK, NFκB, TGFβ, Notch, JAK-STAT1/2, and JAK- STAT3 pathways has been described before, and can in principle be used to analyze pathway activity on any cell type (Bouwman et al., 2020),(Verhaegh et al., 2014),(Stolpe et al., 2019),(Canté-Barrett et al., 2020),(Wesseling-Rozendaal et al., 2021),(van Ooijen et al., 2018) including blood cells (Bouwman et al., 2020),(Wesseling-Rozendaal et al., 2020),(van de Stolpe et al., 2021),(Bouwman et al., 2021). For the current study, quantitative pathway activity scores (PAS) were measured on Affymetrix expression microarray data.

Publicly available Affymetrix expression microarray data were derived from the GEO database (https://www.ncbi.nlm.nih.gov/gds/) (“GEO database: https://www.ncbi.nlm.nih.gov/gds/,” n.d.). For each signaling pathway, PAS were presented on a log2 odds scale. The log2 odds score for pathway activity is derived from the probability score for activity of the pathway-associated transcription factor calculated by the computational model, as described (Verhaegh et al., 2014),(van Ooijen et al., 2018),(Stolpe et al., 2019). Of note, PI3K pathway activity is inversely related to the measured activity of transcription factor FOXO (FOXO PAS) (van Ooijen et al., 2018). For this reason, FOXO activity score is presented in the figures, while interpretation regarding PI3K pathway activity can be found in legends and text.

### Determination of an Immune-signal transduction Pathway Activity Profile (I-PAP)

Combining the PAS for the nine measured STPs we tried to derive characteristic I-PAPs for the analyzed immune cell types and their functional states. Since we did not (yet) want to use absolute PAS for this purpose, relative differences (significant) in PAS between different immune cell types, or between resting/naïve and activated functional state of a specific immune cell type, were used. When an STP was significantly more active in a cell type (e.g. monocytes) compared to another cell type or types, the STP was included in the I-PAP of that cell type (monocytes).

Similarly, when an STP was more active in one functional state (e.g. monocyte, resting state) compared to the other (monocyte activated) the STP was include in the I-PAP of that functional state (monocyte, resting state).

For CD8+ and NK cells, resting/naïve compared to activated functional state, sample numbers were too low (<3) to perform statistics. For these cases we decided to determine tentative I-PAPs by including STPs that had no overlapping PAS scores between the compared functional states.

If two analyzed datasets were available for the same cell type (noted by an asterisk in Table 1-4), allowing comparison of results between independent studies, analysis results from one set are in the Supplement.

**Table 1.**
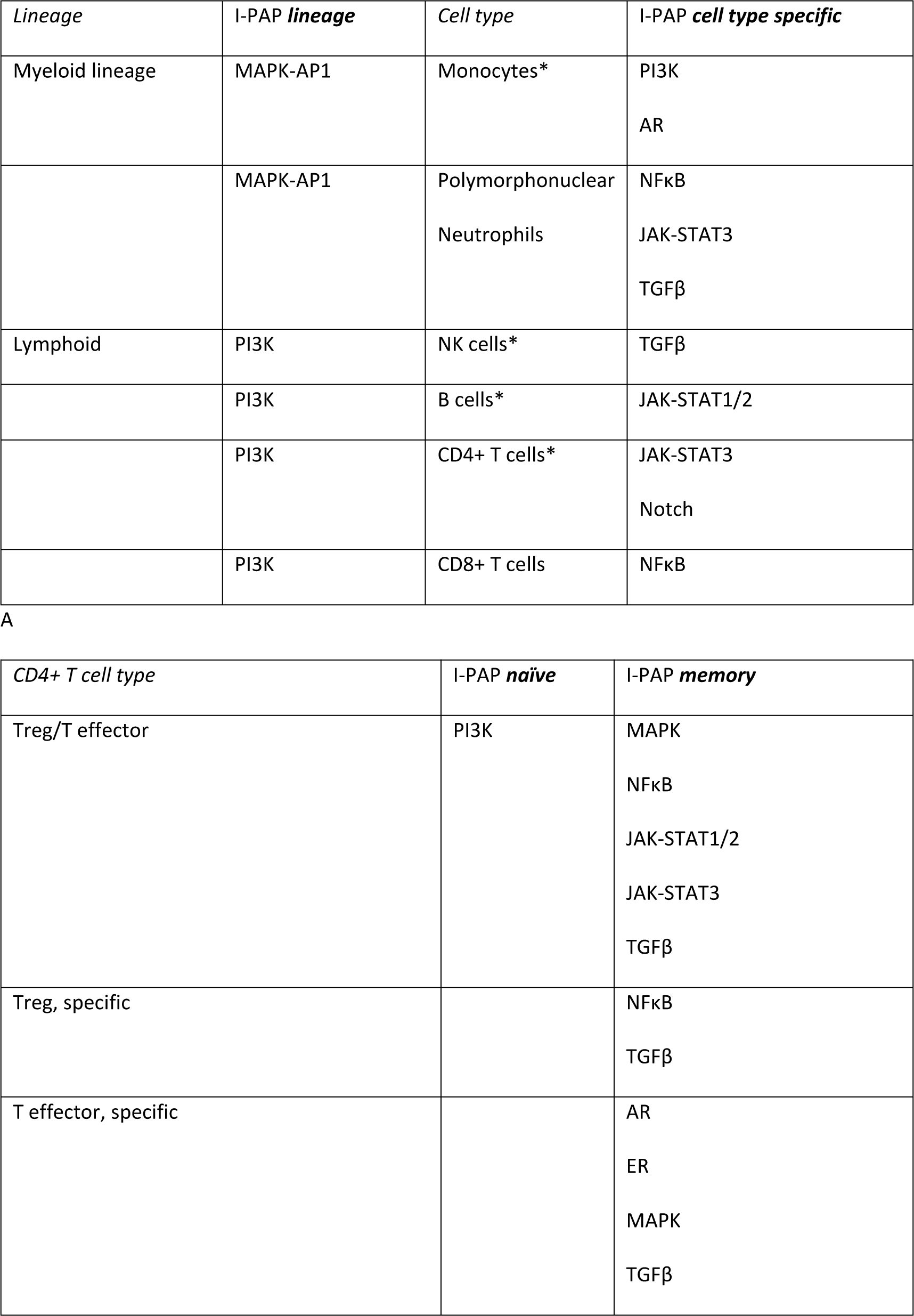
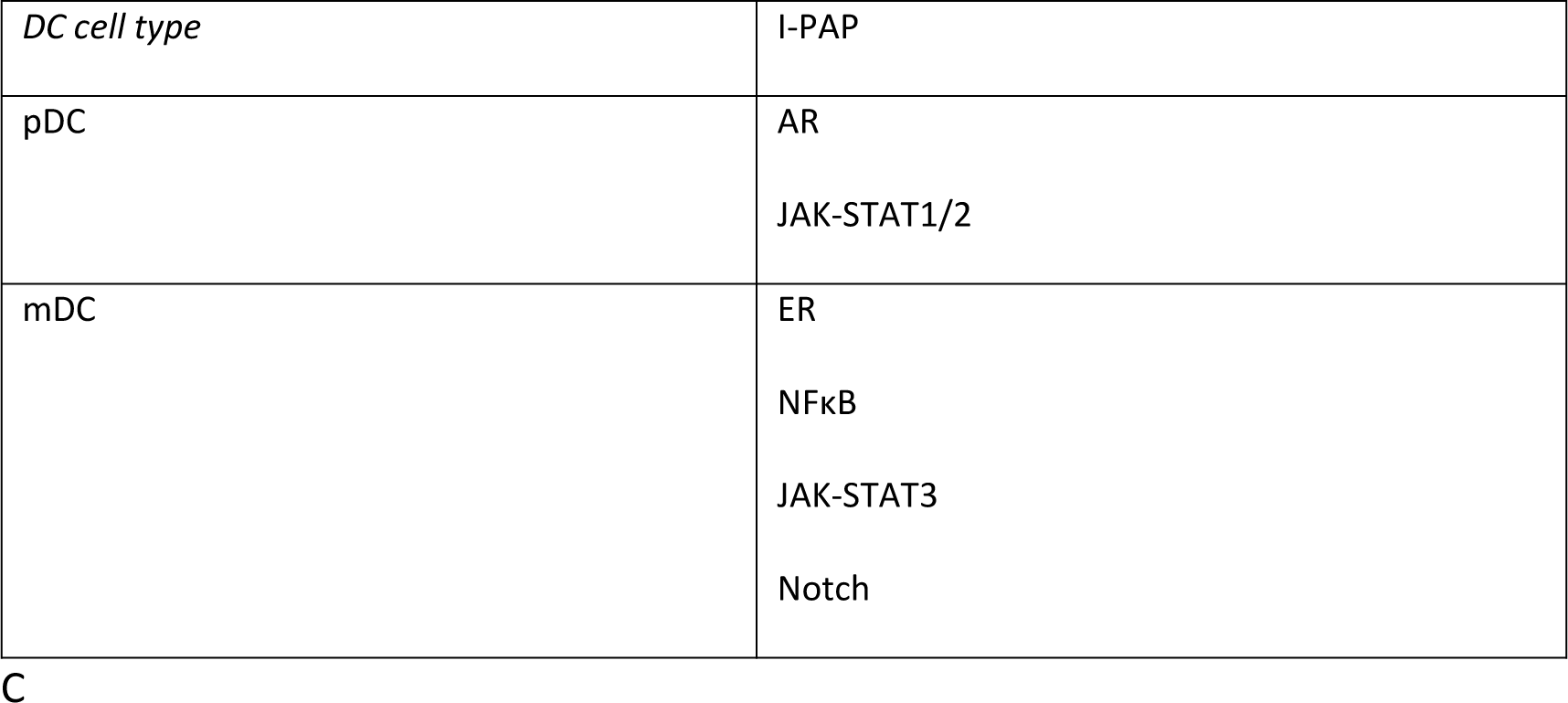
A. I-PAP for myeloid and lymphoid lineage, and I-PAP for cell types within a lineage. I-PAP **lineage**: STPs that are differentially active in lymphoid versus myeloid lineage; I-PAP **cell type specific**: STPs that are differentially active in the indicated cell type within a lineage (in addition to the lineage I-PAP). *B.* I-PAP for CD4+ naïve versus memory T effector and Treg cells combined (Treg/Teffector), and I-PAP for Treg naïve versus Treg memory (Treg specific) and Teffector naïve versus Teffector memory (Treg specific). I-PAP **naïve:** STPs that are more active in both naïve Treg and Teffector cells (Treg/Teffector); or more active in naïve Treg cells specifically (Treg specific) or naïve Teffector cells specifically (Teffector specific); I-PAP **memory:** STPs that are more active in both memory Treg and Teffector cells (Treg/Teffector); or more active in memory Treg cells specifically (Treg specific) or memory Teffector cells specifically (Teffector specific). *C.* I-PAP for myeloid (mDC) and plasmacytoid (pDC) dendritic cells. I-PAP: STPs that are differentially active in mDCs versus pDCs. The pathway that is listed is significantly more active (p<0.01), see Figure 2. * Second independent dataset analyzed, see Supplement.

| *Lineage* | *I-PAP lineage* | *Cell type* | *I-PAP cell type specific* |
| --- | --- | --- | --- |
| Myeloid lineage | MAPK-AP1 | Monocytes* | PI3K<br>AR |
|  | MAPK-AP1 | Polymorphonuclear<br>Neutrophils | NFκB<br>JAK-STAT3<br>TGFβ |
| Lymphoid | PI3K | NK cells* | TGFβ |
|  | PI3K | B cells* | JAK-STAT1/2 |
|  | PI3K | CD4+ T cells* | JAK-STAT3<br>Notch |
|  | PI3K | CD8+ T cells | NFκB |

A
| *CD4+ T cell type* | *I-PAP naïve* | *I-PAP memory* |
| --- | --- | --- |
| Treg/T effector | PI3K | MAPK<br>NFκB<br>JAK-STAT1/2<br>JAK-STAT3<br>TGFβ |
| Treg, specific |  | NFκB<br>TGFβ |
| T effector, specific |  | AR<br>ER<br>MAPK<br>TGFβ |

|  |  |
| --- | --- |
| DC cell type | I-PAP |
| pDC | AR<br>JAK-STAT1/2 |
| mDC | ER<br>NFkB<br>JAK-STAT3<br>Notch |

### Microarray Data Quality Control

Quality control (QC) was performed on Affymetrix data of each individual sample, as described before (Stolpe et al., 2019). In summary, QC parameters include: the average value of all probe intensities, presence of negative, or extremely high (> 16-bit) intensity values, poly-A RNA (sample preparation spike-ins), and labelled cRNA (hybridization spike ins) controls, *GAPDH*, and *ACTB* 3’/5’ ratio, centre of intensity and values of positive and negative border controls determined by affyQCReport package in R, and an RNA degradation value determined by the AffyRNAdeg function from the Affymetrix package in R (Parman C, Halling C, Gentleman R., n.d.), (Gautier et al., 2004). Sample data that failed QC were removed prior to data analysis, except for the neutrophil datasets that failed on reference genes QC parameter, and the B cell dataset that failed on CMoff QC-parameter.

### Affymetrix microarray datasets from preclinical and clinical studies

The following AffymetrixU133Plus2.0 microarray expression datasets (publicly available) from preclinical and clinical studies, deposited in the GEO database, were analyzed (“GEO database: https://www.ncbi.nlm.nih.gov/gds/,” n.d.).

For each sample the type of immune cell (e.g. NK cell) and functional activity (e.g. resting/naïve or activated) was defined by the authors of the corresponding paper or present in the annotation provided by GEO.

1. GSE72642. Immune blood cell types (CD4+ T cells, CD8+ T cells, CD19+ B cells, CD56+ natural killer cells, CD14+ monocytes and PMN (polymorphonuclear cells) isolated from peripheral blood from healthy volunteers (Du et al., 2006).
2. GSE15743. Natural Killer (NK) cells isolated from peripheral blood of healthy individuals, resting and activated by incubation for six hours with two concentrations (1 and 100 ng/ml) interferon alpha (IFNα), *in vitro* (Stegmann et al., 2010).
3. GSE28490. Neutrophils, Eosinophils, Monocytes, CD4+ T cells, CD8+ T cells, NK cells, B cells, pDCs and mDCs isolated from healthy human blood. Cell types were isolated from 5 pools of 5 healthy donors each, with the exception for monocytes, where 10 pools were used*. (Allantaz et al., 2012)*
4. GSE22103. Neutrophils, isolated from peripheral blood of healthy individuals, resting and activated with Escherichia coli lipopolysaccharide (LPS) or granulocyte-macrophage colony-stimulating factor (GM-CSF) and interferon-gamma (INF-g) *in vitro* (Kotz et al., 2010). Note: Sample data from this dataset did not pass our QC criteria. However, RNA Integrity Number (RIN) data were reported to be in the range of 7.4– 9.9, excluding RNA degradation, while comparison with neutrophils from GSE72642 showed a similar STP activity profile; for these reasons the dataset analysis was included.
5. GSE38351. Monocytes, isolated from peripheral blood of healthy individuals, resting and activated for 1.5 hour by incubation with TNFα, IFNα2a and IFNγ *in vitro* (Smiljanovic et al., 2012).
6. GSE43596. Monocytes isolated from peripheral blood of healthy volunteers were differentiated to macrophages, resting and activated by incubation for 2 hours with LPS (100 ng/mL) *in vitro* (Lowe et al., 2014).
7. GSE71566. CD4+ T cells derived from cord blood, resting or activated with anti-CD23/CD28 and differentiated towards respectively T-helper-1 (Th1) (incubation for 72 hrs with 2.5 ng/ml IL12 and Th2 neutralizing antibody anti-IL4, 1 μg/ml) and T-helper-2 (Th2) (10 ng/ml IL4 plus Th1 neutralizing antibody anti-interferon γ, 1 μg/ml) phenotype *in vitro* (Kanduri et al., 2015).
8. GSE63129. CD8+ T cells isolated from healthy individuals, activated by specific antigen, and immune-suppressed by presence of Treg cells *in vitro* (Maeda et al., 2014)
9. GSE39411. B cells, isolated from peripheral blood of healthy individuals, resting and activated for 1 hr and 3 hrs with anti-IgM*, in vitro* (Vallat et al., 2013).
10. GSE18791. Monocytes were isolated from healthy donors and differentiated to dendritic cells. Dendritic cells were activated by infection with Newcastle disease virus (NDV) *in vitro* (Zaslavsky et al., 2010).
11. GSE65010. Clinical study. Treg and Teffector cells (naïve and memory), isolated from peripheral blood of healthy individuals and RA patients (Walter et al., 2016)
12. GSE93272. Clinical study. Whole blood samples from healthy donors and RA patients. (Tasaki et al., 2018). Only Sample data from patients without treatment were used.

### Privacy

All analyzed data were obtained in agreement with GDPR regulations, as implemented within Philips.

### General rules for interpretation of signal transduction pathway activity score (PAS)

An important and unique advantage of the pathway activity assays is that in principle they can be performed on each cell type. A few considerations to bear in mind when interpretating log2 odds PAS, as described before (van de Stolpe et al., 2019):

1. On the same sample, log2 odds PAS cannot be compared between different STPs, since each STP has its own range in log2 odds activity scores (van de Stolpe et al., 2019)
2. The log2 odds range for STP activity (minimum to maximum activity) may vary depending on cell type. Once the range has been defined using samples with known STP activity, on each new sample the absolute value can be directly interpreted against that reference. If the range has not been defined, only differences in log2 odds activity score between samples should be interpreted.
3. PAS are highly quantitative, and even small differences in log2 odds PAS can be reproducible and meaningful.
4. A negative log2 odds ratio does not necessarily mean that the STP is inactive.

### Statistics

T-tests (unpaired, two-sided) were performed to compare pathway activity scores across groups. In case another statistical method was more appropriate due to the content of a specific dataset, this is indicated in the legend of the respective figure. Pearson correlation tests were performed.

Exact p-values are indicated in the figures. P<0.01 was considered significant (Amrhein et al., 2019). For studies with sample numbers n<3, descriptive analysis was performed and no I-PAP was included in the tables (NK cells, CD8+ T cells). For Tables 1-4 (I-PAPs), STPs were included for which the PAS difference was statistically significant. Underneath figures tables are shown in which numbered sample groups are compared using a t-test, deltas indicate the absolute measured difference in PAS between the compared sample groups, for every STP.

## Results

The immune response is determined by controlled activity of immune cells that are part of the innate and adaptive immune system, with the dendritic cell as the most powerful antigen-presenting cell bridging the two parts of the immune system (Figure 1A) (Davis, n.d.).

To define I-PAPs associated with the functional activity state of these immune cell types, STP activity analysis (AR, ER, PI3K-FOXO, MAPK, JAK-STAT1/2, JAK-STAT3, NFκB, TGFβ, Notch pathways) was performed on data from studies in which the respective immune cells were derived from healthy donors and activated *in vitro* in a controlled manner.

### STP analysis of primary immune cells isolated form blood

*Naïve/Resting state I-PAP of Monocyte, polymorphonuclear neutrophils (PMN), B cells, CD4+ and CD8+ T cells, NK cells* (Table 1A, Figure 2A) Signaling pathway activities of resting immune cells were more similar between different healthy individuals (n=3) than between the different blood cell types. Myeloid (PMN, monocytes) and lymphoid (B and T cells) blood cell lineages could be easily distinguished; within the two lineages I-PAP profiles were more similar than between the lineages. Compared to lymphoid cells, myeloid cell types (monocytes and polymorphonuclear neutrophils) had higher FOXO PAS, reflecting lower activity of the PI3K growth factor pathway, and higher MAPK-AP1 and NFκB PAS; lymphoid cell types showed higher PI3K pathway activity (lower FOXO PAS) (Figure 2A).

**Figure 2.**
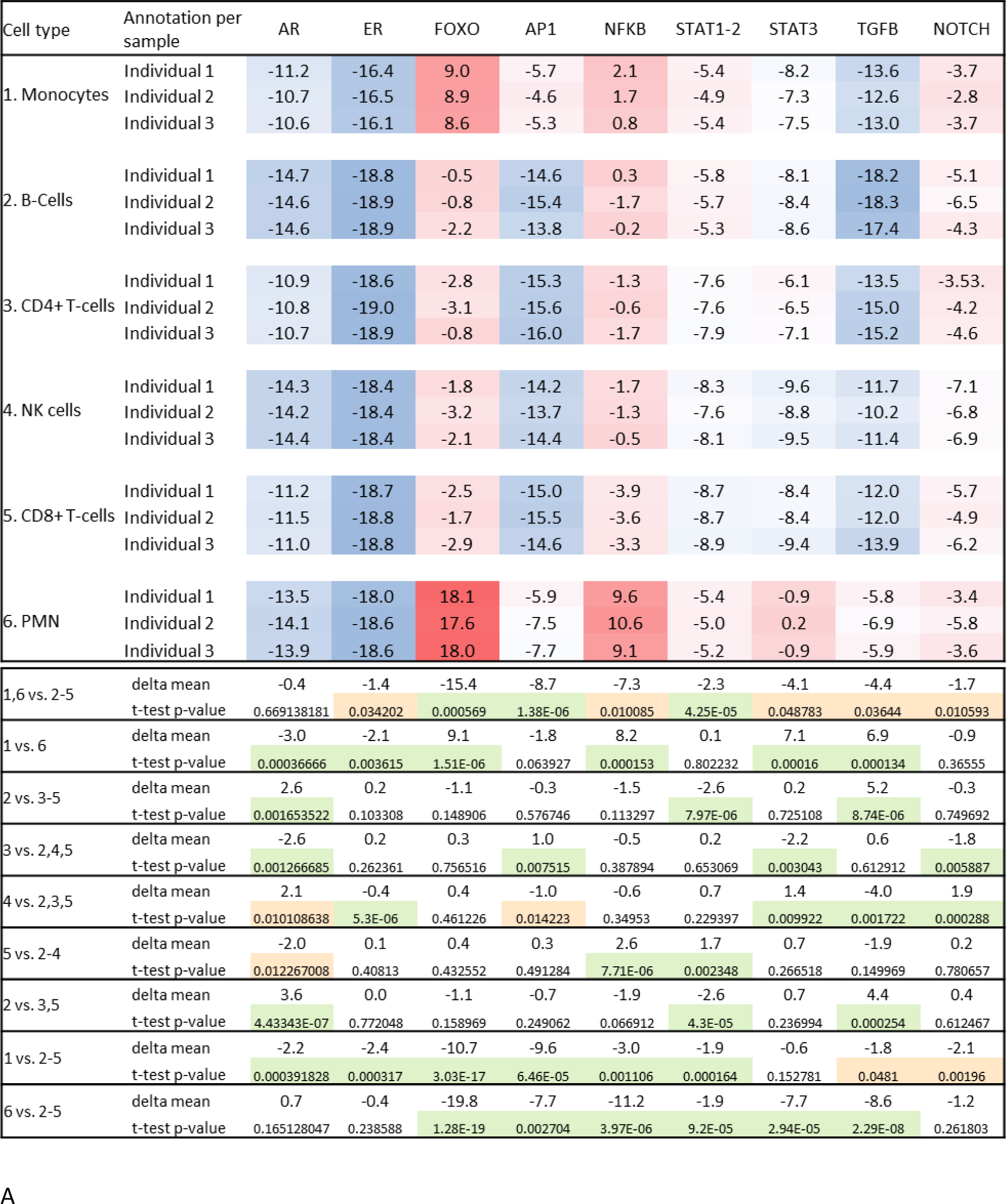
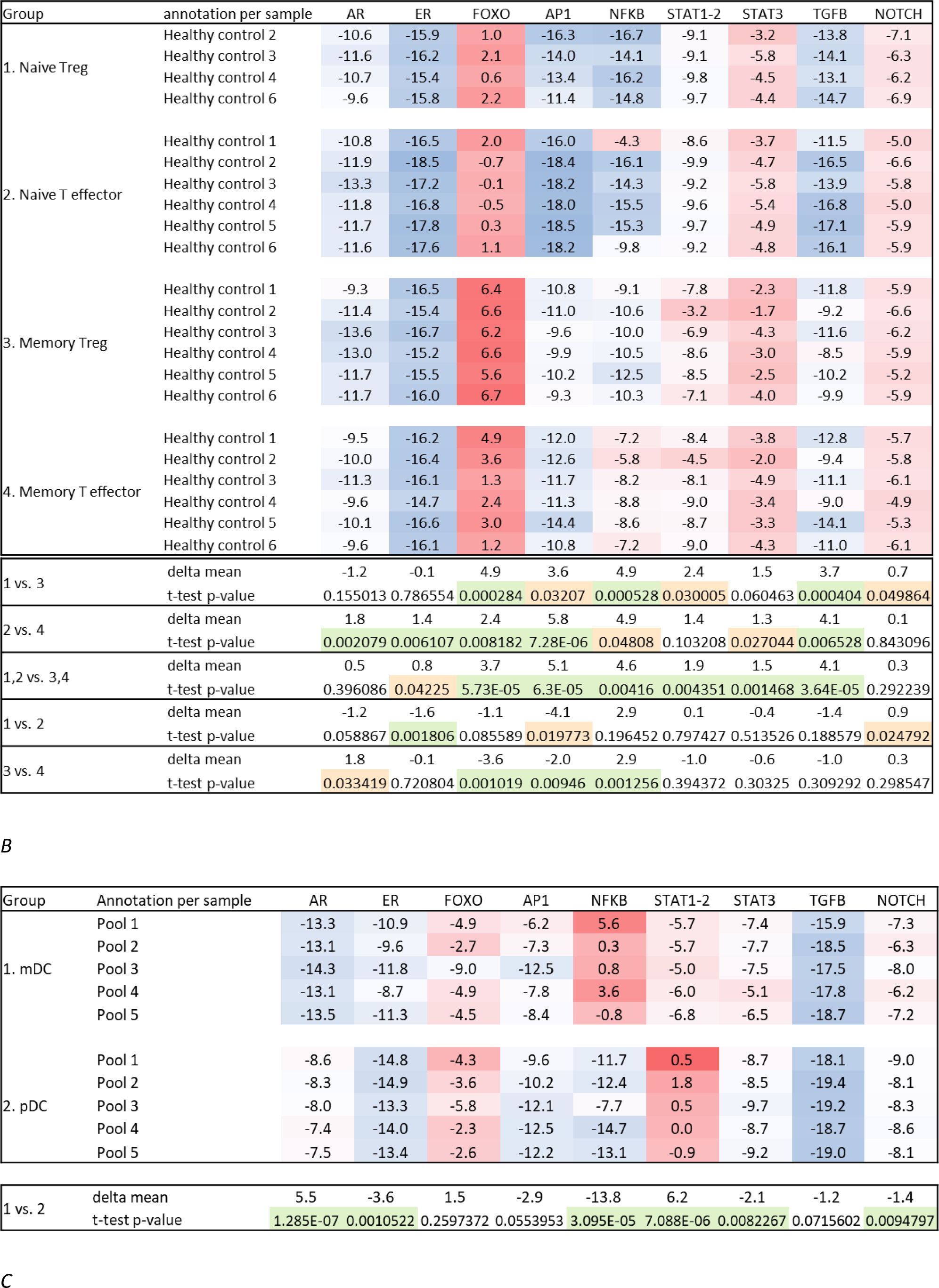
A. GSE72642 (Du et al., 2006). Signal transduction pathway activity for different naïve immune cell types isolated from blood of 3 healthy donors. B. GSE65010 (Walter et al., 2016). Naïve and memory CD4+ Treg and T effector cells isolated from blood samples of healthy volunteers. C. GSE28490 (Allantaz et al., 2012). Myeloid (mDC) and plasmacytoid (pDC) subsets of dendritic cells isolated from 5 pooled blood samples, each 5 healthy volunteers. Top: PAS for the respective signal transduction pathways per individual analyzed sample. First column contains individual sample annotation; top of column contains name of signaling pathway; FOXO transcription factor activity is the inverse of PI3K pathway activity. Signaling pathway activity scores are presented on log2 odds scale; color coding ranging from blue (most inactive) to red (most active). For cell isolation protocols see dataset-associated publication. Bottom: Statistical comparisons between sample groups designated by numbers: exact p-values and the delta in PAS are provided. P<0.01, green; p<0.5, orange.

Within the lineages, cell types showed variations in STP activity, defining their I-PAP (Figure 2A). For the lymphoid lineage (high PI3K pathway activity) B cells had highest JAK-STAT1/2 PAS, CD4+ T cells highest JAK-STAT3 PAS, NK cells by highest TGFβ PAS, and CD8+ T cells lowest NFκB PAS.

For the myeloid lineage (high MAPK, NFκB pathwya activity), neutrophils has higher NFκB, JAK- STAT3, and TGFβ PAS than monocytes (Figure 2A).

### Naïve and memory CD4+ T cells (Figure 1B, Table 1B)

T cells change into memory T cells upon disappearance of the antigen, but they retain the memory to the antigen, enabling rapid activation upon renewed exposure to the specific antigen.

Memory CD4+ T cells (both Treg and T effector) had lower PI3K pathway activity than naïve T cells, reflecting loss of division, and higher PAS for a number of STPs, including the TGFβ pathway (Table 1B, Treg/ Teffector).

While having an allmost identical I-PAP in the naïve state, memory Treg and Teffector cells could be clearly distinguished by higher NFκB and TGFβ PAS in Tregs (Table 1B, Treg specific) and higher AR, ER and MAPK PAS in Teffector cells (Table 1B, T effector specific).

*Dendritic cells (Table 1C, Figure 2C)*.

Dendritic cells function as master regulators of the immune response. Blocking functional activation results in immunosuppression, as for example seen in cancer and certain viral infections. Plasmacytoid (pDC) and myeloid (mDC) DC subsets recognize different pathogens and activate innate and adaptive immune response in a specific manner. pDC precursors produce type I interferons in response to viruses and can activate mDCs.

In line with their different functions, plasmacytoid and myeloid dendritic cells (pDCs and mDCs) had different I-PAPs, with most prominent differences in the NFκB, AR, and JAK-STAT1/2 PAS.

### STP analysis of primary immune cells from blood, activated in vitro

Following STP analysis of immune cells isolated from blood, we investigated changes in signaling pathway activity upon *in vitro* activation of immune cells.

### Cells of the innate immune response, Natural Killer cells

NK cells are part of the lymploid lineage but function in the innate immune response as cyotoxic cells that kill cells that do not express the correct Major Histocompatibility Complex (MHC) signature.

Primary NK cells had been activated *in vitro* with both a low and high dose interferon α. The low sample number only allows a descriptive analysis. Compared to control incubation, activation resulted dose-dependently in reduced PI3K (increased FOXO PAS), NFκB and Notch pathway activity, and increased JAK-STAT pathway activity (JAK-STAT1/2 and JAK-STAT3 PAS) (Figure 3A).

**Figure 3.**
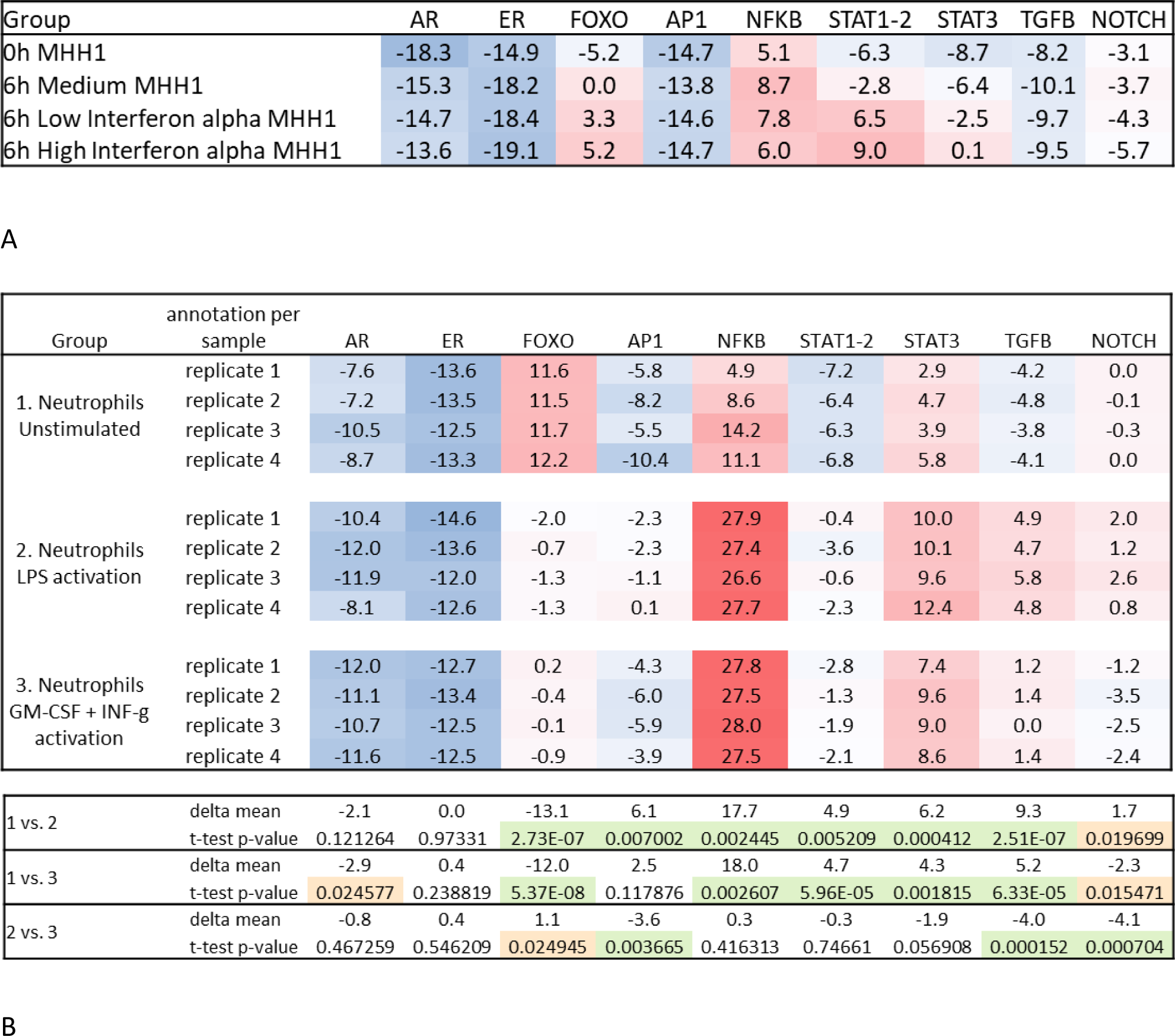
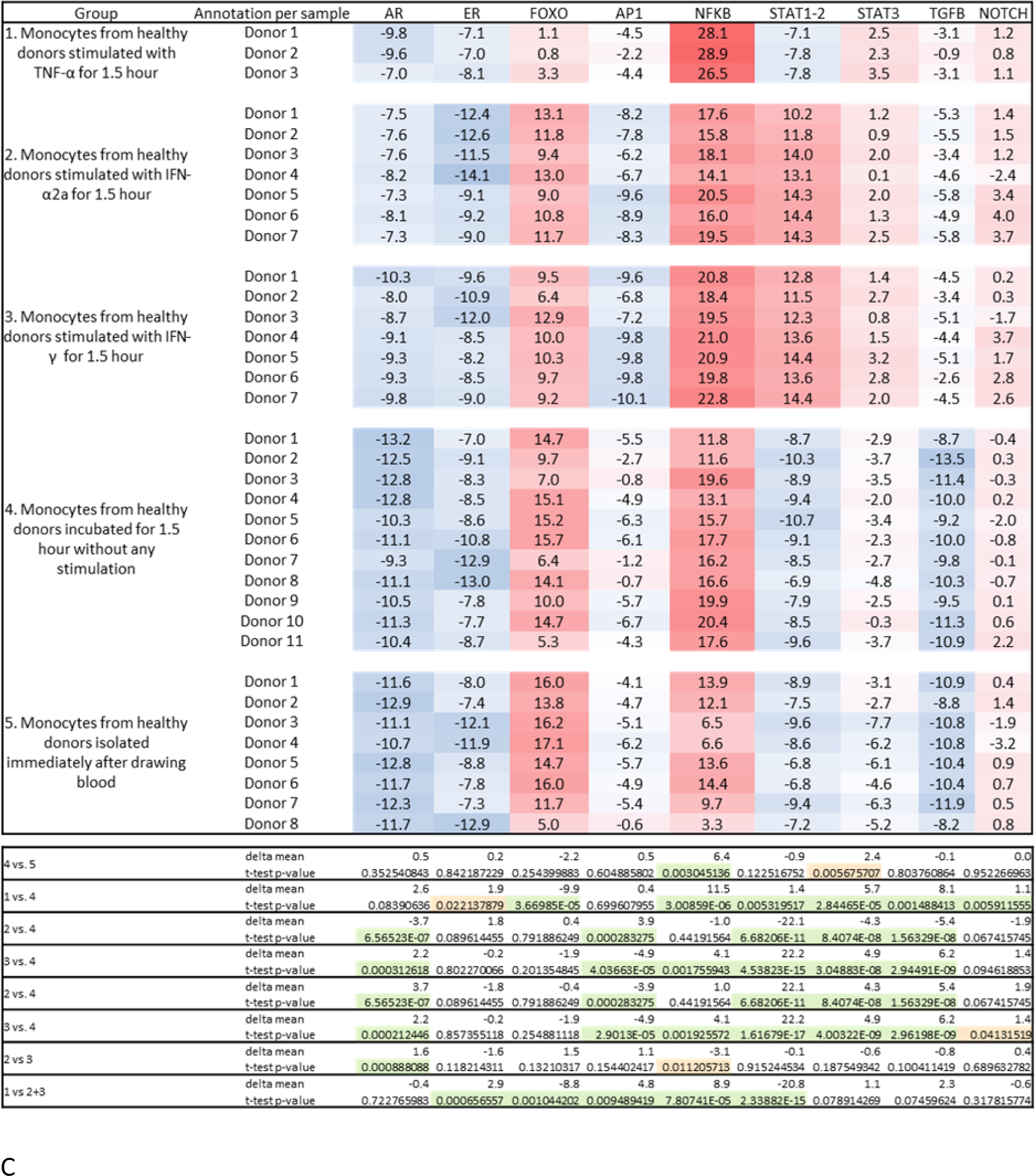
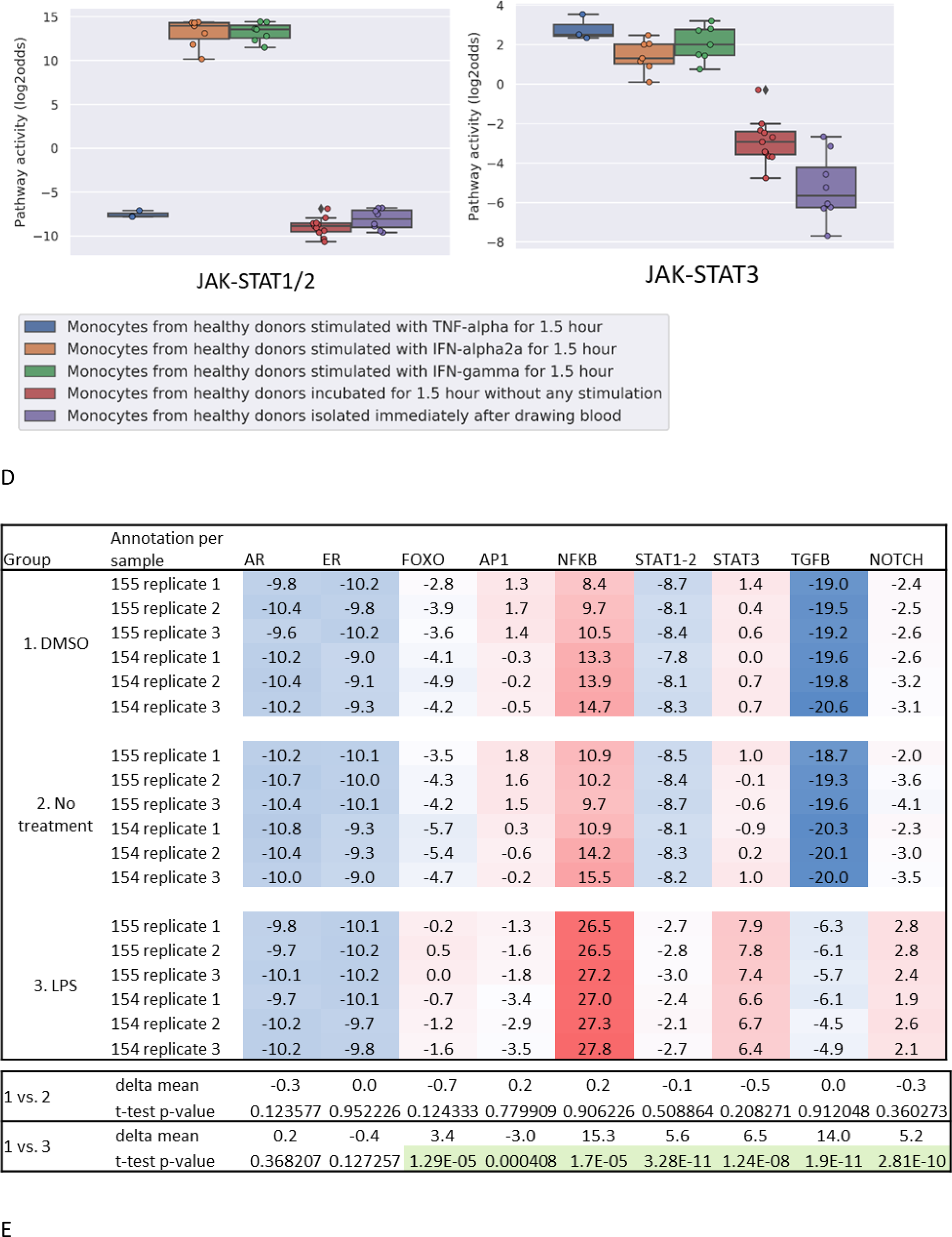
Immune cell types from the innate immune system, effect of immune cell activation. A. GSE15743 (Stegmann et al., 2010). Human NK cells, resting and activated by treatment with IFNα for 6 hrs. B. GSE22103 (Kotz et al., 2010); Neutrophils, resting and activated in vitro in whole blood with Escherichia coli lipopolysaccharide (LPS), with granulocyte-macrophage colony-simulating factor (GM-CSF) and interferon-gamma (INF-g) (GM+I) (in a microfluidic cassette). Sample data from this dataset did not pass our (rigorous) QC criteria. C. GSE38351 (Smiljanovic et al., 2012). Monocytes of healthy volunteers, in vitro activated with TNFα, IFNα2a or IFNγ for 1.5 hour. JAK-STAT1/2 PAS have been reported before, Bouwman et al, doi: 10.3389/fimmu.2020.575074. D. GSE38351 (Smiljanovic et al., 2012). JAK-STAT1/2 and JAK-STAT3 pathway activity in monocytes illustrating that IFN type I and type II strongly activate the activate JAK-STAT1/2 pathway in contrast to TNFα; both TNFα and interferons activate the JAK-STAT3 pathway to the same extent. The line inside the rectangle indicates the median of data.The rectangle shows the interquartile range (IQR). Its lower edge is placed at the 25% percentile (1st quartile). The upper edge is at the 75% percentile (3rd quartile). The T-shaped lines are the whiskers. Normally the range of the whiskers shows values which are between the 1st quartile (Q1) and a number (Q1 — IQR1.5). The upper whisker ends at the value = Q3 + IQR1.5. JAK-STAT1/2 PAS have been reported before, Bouwman et al, doi: 10.3389/fimmu.2020.575074. E. GSE43596 (Lowe et al., 2014). Macrophages, from two donors (154 and 155) measured in triplicate, resting (no treatment and DMSO vehicle) and in vitro activated with LPS.

### Cells of the innate immune response, Neutrophils (Figure 3B, Table 2)

Neutrophils phagocytose pathogens and mediate important inflammatory reactions within the innate immune response.

**Table 2.** I-PAP of in vitro cytokine-activated immune cells of the innate immune system and dendritic cells, compared to their resting immune counterpart. I-PAP **resting**: STPs that are more active in functionally resting/naïve cells; I-PAP **activated**: STPs that are more active in functionally activated cells. Shared activated pathways: STPs that are activated by multiple used stimuli; Specific activated pathways: STPs that are specifically activated by the stimulus (in addition to shared activated STPs). The pathway that is listed is significantly more active (p<0.01), see Figure 3 and 5. NDV: Newcastle Disease Virus. * other independent dataset(s) analyzed, see Supplement

| Cell type,<br>*activator used* | I-PAP **resting** | I-PAP **activated cells** |
| --- | --- | --- |
| Neutrophils,<br>*activated by LPS, GM-CSF, IFN (shared activated pathways)* |  | PI3K<br>NFκB<br>JAK-STAT1/2<br>JAK-STAT3<br>TGFβ |
| Neutrophils,<br>*LPS, specific activated pathways* |  | MAPK |
| Monocytes*,<br>*activated by TNFα, IFNα, or IFNγ (shared activated pathways)* |  | JAK-STAT3<br>TGFβ<br>JAK-STAT1/2 |
| Monocytes,<br>*TNFα, specific activated pathways* |  | PI3K<br>NFκB |
| Monocytes,<br>*Interferon, specific activated pathways* |  | AR<br>MAPK<br>JAK-STAT1/2<br>Notch |
| Macrophages*,<br>*activated by LPS* | PI3K<br>MAPK | NFκB<br>JAK-STAT1/2<br>JAK-STAT3<br>TGFβ<br>Notch |
| Dendritic cells*,<br>*activated by infection with NDV (virus)* |  | NFκB<br>JAK-STAT1/2<br>TGFβ |

Activation of neutrophils with LPS or with granulocyte-macrophage colony-stimulating factor (GM-CSF) plus interferon-gamma (INF-γ) resulted in activation of a number of shared STPs (Table 2, *I-PAP activated cells*). In addition to these changes, LPS specifically induced MAPK pathway activity. Of all investigated immune cell types, activated neutrophils had highest TGFβ PAS.

### Cells of the innate immune response, Monocytes (Figure 3C, 3D, Table 2)

Monocytes are the circulating precursor cells of tissue macrophages, and home to tissue locations to infiltrate and differentiate to tissue macrophages. Monocyte activation-induced JAK- STAT1/2 pathway activity PAS have been reported before and have been included here to enable determination of the complete monocyte I-PAP (Bouwman et al., 2020).

Brief *in vitro* incubation itself induced a small increase in NFκB and JAK-STAT3 PAS. Compared with this control state, activation with TNFα, interferon type I (IFN-α2a), or interferon type II (IFN-γ) induced activity of shared STPs (*Table 2, monocytes, shared activated pathways*), while in addition STPs were specifically activated by TNFα and the two types of interferon (Table 2, TNFα, specific activated pathways; interferon, specific activated pathways). While activation with TNFα or either type of interferon resulted in a comparable induction of JAK-STAT3 PAS, the large increase in JAK-STAT1/2 PAS was specific for activation with interferons (delta in log 2 odds: 1.4 for TNFα and 22 for interferons), (Figure 3D). Interferons are specific activators of the JAK- STAT1/2 pathway (Platanias, 2005)

### Cells of the innate immune response, Macrophages (Figure 3E) (Table 2)

Activation with LPS resulted in a reduction in growth factor pathway activity (both PI3K and MAPK pathways) and increased NFκB, JAK-STAT3, TGFβ, and Notch PAS. This I-PAP was comparable with the I-PAP of LPS-stimulated neutrophils, except for higher TGFβ PAS in neutrophils (compare Figure 3E with 3B).

### Cells of the adaptive immune response, CD4+ T cells

CD4+ T cells regulate activity of CD8+ T cells and B cells and can differentiate towards specific T helper cells. The Th1 type supports the cell-mediated immune response and helps cytotoxic T cells while Th2 cells mainly help B cells in their humoral (antibody) immune response.

Except for ER, AR, and MAPK PAS, pathway analysis results of CD4+ T cells and Th1 and Th2 cells have been reported before, and are presented to enable determination of a complete I-PAP (Wesseling-Rozendaal et al., 2020)(Figure 4A, Table 3). Differentiation of activated CD4+ T cells to lymphocyte subtypes Th1 and Th2 resulted in opposite changes in NFκB PAS: increased NFκB pathway activity in Th1, and decreased activity in Th2. Incubation of activated CD4+ T cells with an antibody against interferon-γ was used to induce Th2 cell differentiation *in vitro*, and this resulted in the expected reduction in PAS for the interferon-inducible JAK-STAT1/2 pathway.

**Figure 4.**
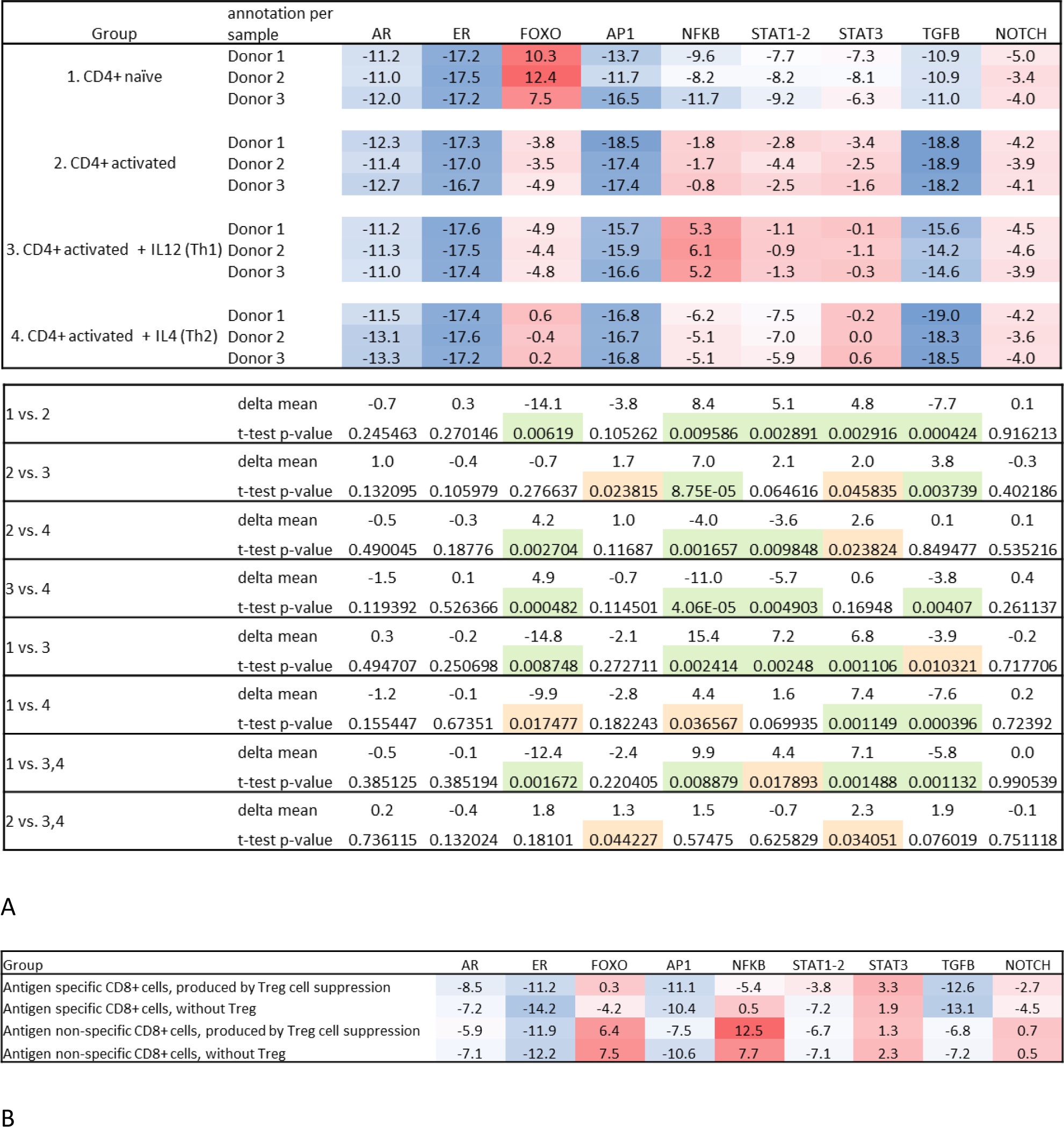
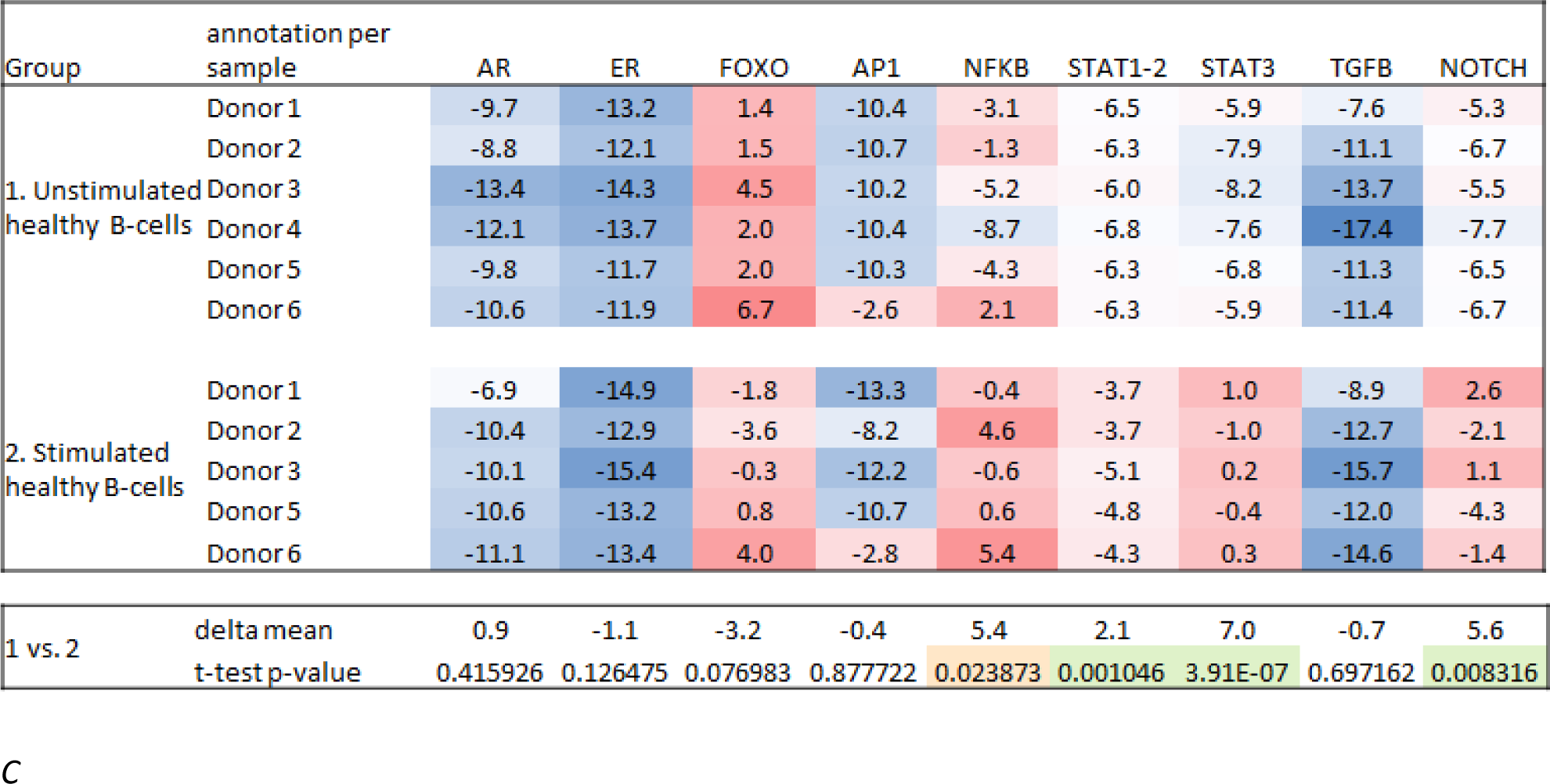
Immune cell types from the adaptive immune system, effect of immune activation. A. GSE71566, CD4+ T cells derived from cord blood of 3 donors, resting or activated using anti CD3/CD28 (A), or stimulated with IL-12 and IL-4 for differentiation towards respectively T-helper-1 (Th1) and T-helper-2 (Th2) T cell phenotype. (partly reproduced from: Wesseling et al, https://www.biorxiv.org/content/10.1101/2020.10.08.292557v1, currently under review Frontiers Immunology) B. GSE63129; CD8+ T cells, activated by specific antigen, and immune-suppressed by presence of Treg cells. Antigen-specific (Tet+) and non-specific (Tet-) CD8+ T cells with Treg cells (Treg+) or without Treg (Treg-) (Maeda et al., 2014). C. GSE39411; B cells from healthy donors, resting and activated with anti-IgM for 210 min (Vallat et al., 2013).

**Table 3.**
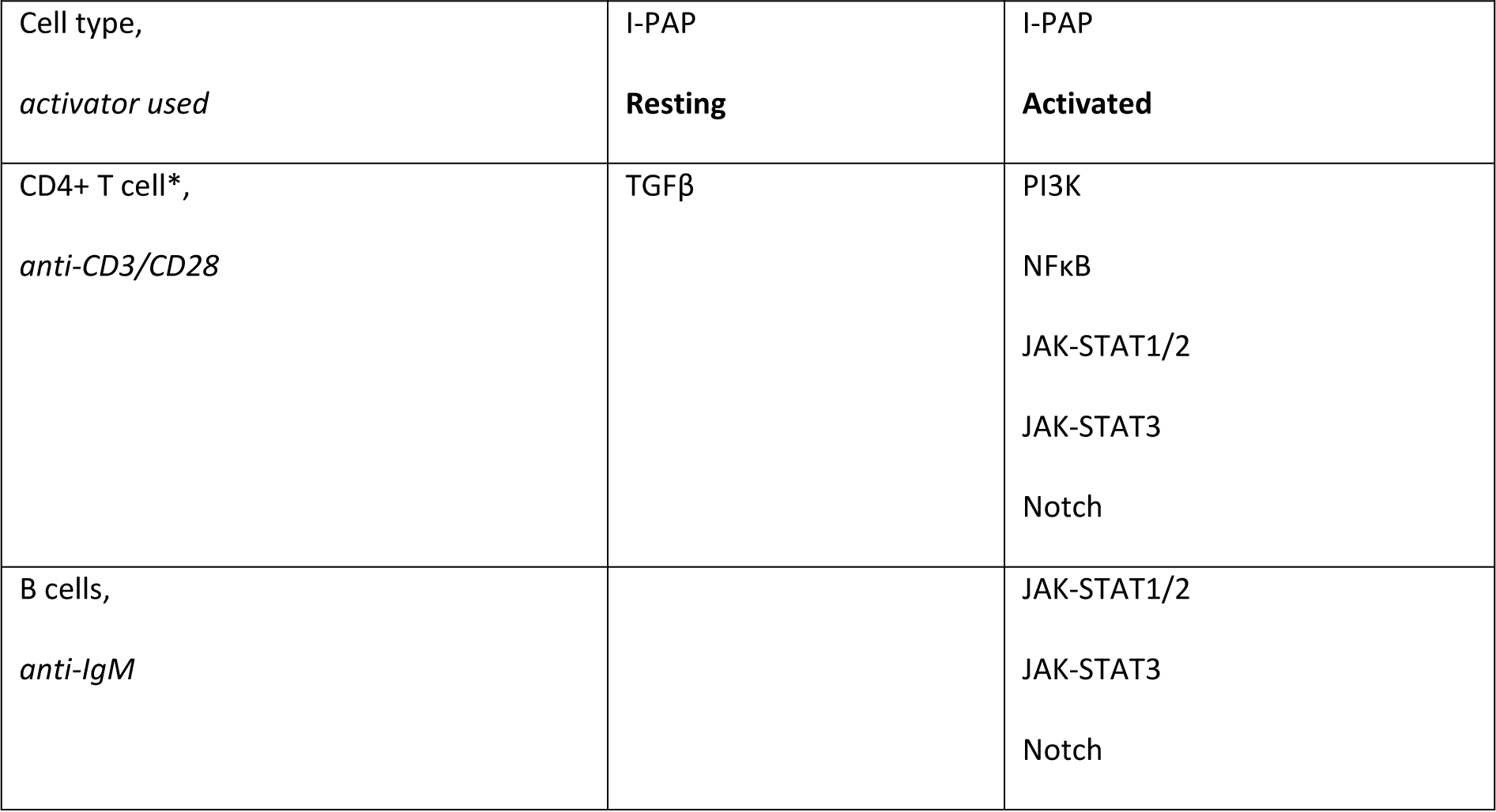
I-PAP of in vitro cytokine-activated immune cells of the adaptive immune system, compared to their resting counterpart. I-PAP **resting**: STPs that are more active in functionally resting/naïve cells; I-PAP **activated**: STPs that are more active in functionally activated cells. The pathway that is listed is significantly more active (p<0.01), see figure 4. * other independent dataset analyzed, see Supplement

*Cells of the adaptive immune response, CD8+ T cells (*n<3, descriptive interpretation, Figure 4B) CD8+ T cells are cytotoxic T cells that attack cells upon specific antigen recognition.

For specific antigen-induced immune-activated CD8+ T cells, data from one healthy individual were available, allowing only a tentative and descriptive I-PAP. Antigen-specific activation (compared to non- antigen-specific activation) was associated with higher PI3K (lower FOXO PAS) pathway activity, reflecting T cell proliferation, and lower NFκB, TGFβ, and Notch PAS; in the presence of immunosuppressive Treg cells, PI3K PAS was lower and NFκB PAS decreased further.

### Cells of the adaptive immune response, B cells (Figure 4C, Table 3)

B cell produce IgM and IgG antibodies directed towards specific antigens. Activation of B cells with IgM antibody resulted in an I-PAP with a notably increased Notch PAS, only equalled by LPS-induced Notch PAS in neutrophils and macrophages, and increased JAK-STAT pathway activity (both JAK-STAT1/2 and JAK-STAT3). (Table 3, Figure 4D).

### ***Dendritic cells*** (Figure 5, Table 2)

*In vitro* infection with the Newcastle Disease virus has been developed as a model system for *in vivo* DC maturation (Zaslavsky et al., 2010). Within two hours after viral infection NFκB PAS increased, followed at four hours by an increase in JAK-STAT1/2 PAS and after 10 hours by moderately increased TGFβ PAS (p=0.01). This is a logical orde of pathway activation since the NFκB pathway is required for antigen processing; NFκB-induced production of interferons is responsible for activation of the JAK-STAT1/2 pathway, necessary for antigen presentation (Barber, 2015); the final activation of the TGFβ pathway probably enables migration of the antigen-presenting DCs to the lymph node (Batlle and Massagué, 2019).

**Figure 5.**
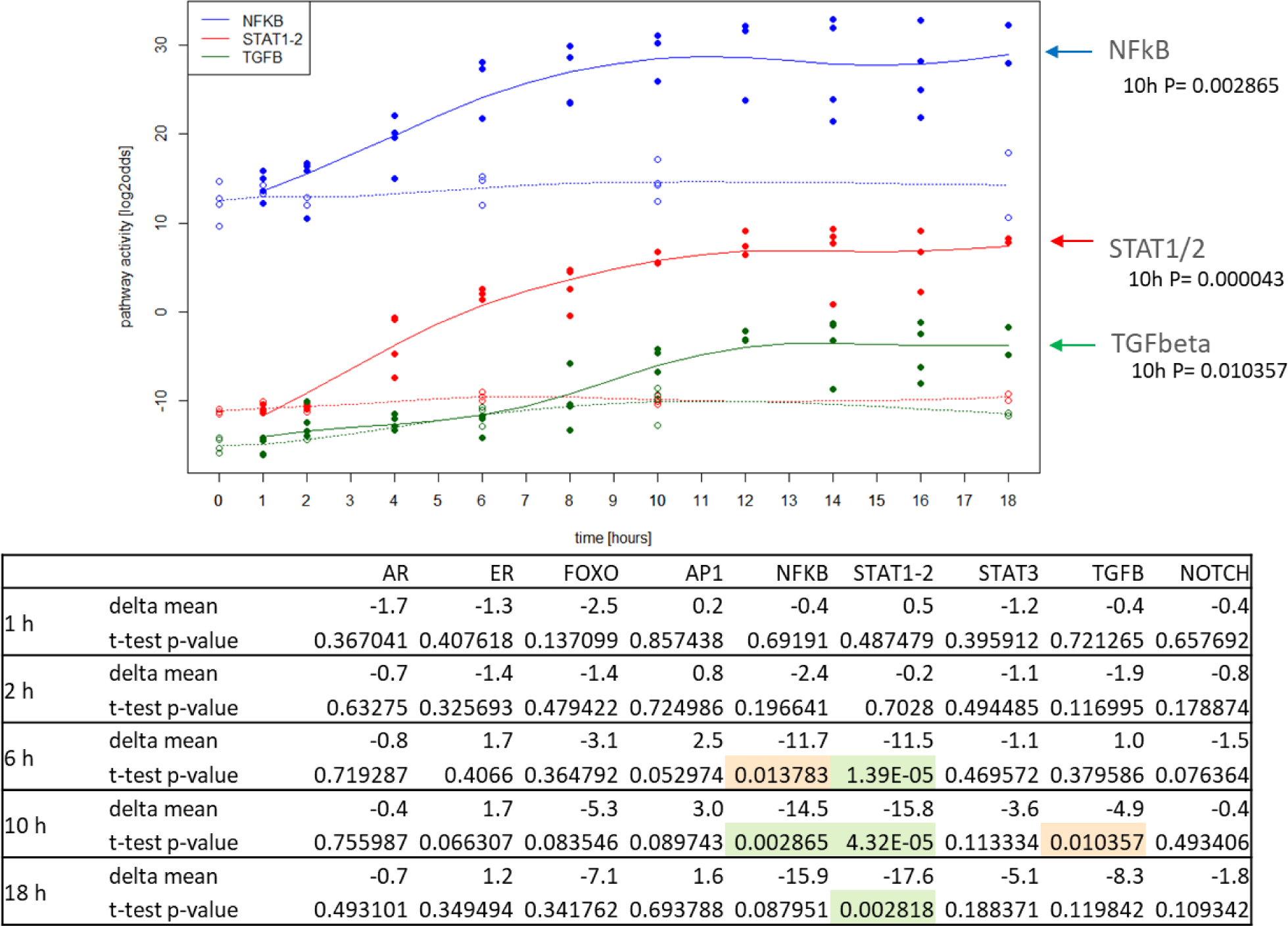
Dendritic cells, effect of immune activation. GSE18791(Zaslavsky et al., 2010). Dendritic cells (Monocyte-derived DCs) from two donors, resting and activated by infection with Newcastle disease virus (NDV), at 1, 2, 4, 6, 8, 10, 12, 14, 16 and 18 hours after infection versus control.

## Reproducibility of STP PAS

### Comparison of pathway activity scores between samples from different healthy donors

In the datasets in which cells were obtained from multiple healthy donors, there was a remarkable consistency in STP PAS between individual donors, both for resting and activated immune cell types, suggesting tight control of functional processes in the immune response (Figure 2-5).

### Comparison of I-PAP of freshly isolated immune cells with cells incubated in vitro

The absolute pathway PAS of immune cells kept for a number of hours *in vitro* differed from the I-PAP identified for primary cells directly analyzed after isolation from peripheral blood, probably under the influence of variations in the composition of the incubation medium (e.g. concentration of growth factors that activate the PI3K pathway) or incubation time, (compare Figure 3,4 with Figure 2 and S1).

### Reproducibility of I-PAPs across independent studies

To confirm the I-PAPs for the various immune cell types, we tried to analyze an independent second dataset for the same cell type (Figures S2-5 in the supplement). This was not possible for all immune cell types (lack of available studies with the required sample data or in case of neutrophil datasets due to QC failure for Affymetrix data). Overall, I-PAPs were comparable between independent datasets and similar relative differences in pathway activities between resting and activated cells were found, despite studies having been performed in different laboratories under different conditions (compare figures 2-5 with S1-S6). However, I-PAP differences were found between independent datasets for a specific immune cell type which were probably caused by different activation protocols, for example, NK cells were either stimulated with IFNα for six hours and compared with control culture medium for the same time period (Figure 3A), or stimulated with IL2 in culture medium for 24 hours without an incubation-time control sample, or with a coctail of IL-2, IL-12, IL-18 for 24 hours (Figure S2).

Analysis of the various immune cell types in a resting state, directly from peripheral blood samples, also showed consistent results between two independent datasets (compare Figure 2 with Figure S2). Besides illustrating reproducibility of the STP assay technology, this provides consistent evidence for the roles of signaling pathways in controlling immune cell functions.

### Summary on I-PAPs of lymphocyte and monocytic lineage cell types in resting versus immune-activated functional state

With respect to growth factor pathways MAPK and PI3K (responsible for cell division), comparing immune cells from the myeloid (neutrophils, monocytes, macrophages, and dendritic cells) with the lymphoid lineage (NK cells; CD4+ T cells, CD8+ T cells, B cells) in a resting state, suggests the MAPK pathway as the dominant growth factor pathway in myeloid-derived cells, and the PI3K pathway in lymphoid cells (Table 1). Between lineage subtypes additional variations in activity of these growth factor pathways was found, e.g. myeloid derived DCs had a higher MAPK pathway activity than their plasmacytoid counterpart (Table 1C) and memory Treg cells typically had lower PI3K PAS scores compared to their naïve counterpart (Table 1B).

In line with a (cell division) role for the PI3K pathway in lymphoid lineage cells, with the exception of NK cells, immune activation was associated with PI3K pathway activation in CD4+ T cells, CD8+ T cells, and B cells (unpaired t-test not significant, paired t-test two sided, p=0.009343), and PI3K pathway activity was also higher in naïve compared to memory (CD4+) T cells (Table 4, Table 1). In myeloid cells growth factor pathway activity did not change upon cell activation.

**Table 4.** I-PAP of CD4+ T cells differentiated to Th phenotype. I-PAP **activated CD4+ Tcell**: STPs that are more active in activated CD4+ T cells compared to their differentiated counterpart. I-PAP **Differentiated to Th cell type** STPs that are more active Th cell phenotypes compared to activated CD4+ T cells. The pathway that is listed is significantly more active (p<0.01). * other independent dataset analyzed, see Supplement

| CD4+ T cell subtype,<br>*cytokine used to differentiate to subtype* | I-PAP<br>**activated CD4+ T cell** | I-PAP<br>**Differentiated to Th**<br>**cell type** |
| --- | --- | --- |
| Th1*,<br>*IL-12* |  | NFκB<br>TGF-β |
| Th2*,<br>*IL-4* | PI3K<br>NFκB<br>JAK-STAT1/2 |  |

TGFβ pathway activity was inversely related to immune activation in lymphoid cell types, but associated with immune activation in myeloid cell types. Immune-activation of CD4+ T cells (and CD8+ T cells, n<3) was associated with reduced TGFβ pathway activity and memory T cells had higher TGFβ PAS than naïve T cells (Table 1). This role of the TGFβ pathway was in line with our earlier observations in activated CD4+ T cells that became tolerogenic upon exposure to breast cancer supernatant, associated with a marked increase in TGFβ pathway activity (Wesseling-Rozendaal et al., 2020). In contrast, in cells of the myeloid lineage, immune activation was associated with increased TGFβ PAS, to much higher values than in lymphoid cells (Table 2,3, Figures 3,4).

NFκB PAS scores were higher in myeloid cells than lymphoid cells, and increased strongly upon activation with highest absolute scores in neutrophils.

The JAK-STAT pathways seemed to play a role in activation of both myeloid and lymphoid cell types, although the JAK-STAT3 pathway activity seemed more prominent in myeloid cells. (Table 2,3).

For many signaling pathways, including NFκB and TGFβ, activation was variable across the different cell types and their functional state, suggesting different functions for these pathways, depending on cellular context.

### Exploration of clinical value: STP analysis of a clinical study on RA

Quantifying the immune response state based on blood sampling may find clinical utility in predicting and monitoring response to signaling pathway-targeted therapy (such as TNFα and JAK inhibitors) in auto-immune diseases like rheumatoid arthritis, among other applications.

We analyzed two clinical studies on rheumatoid arthritis (RA) for which blood sample data were available (Walter et al., 2016),(Tasaki et al., 2018). In whole blood samples TGFβ PAS was increased in both male and female RA patients, while increased JAK-STAT3 PAS reached significance in the female patient group. Many immune cell types may contribute to increased TGFβ and JAK-STAT3 PAS measured in whole blood. The second study allowed analysis of CD4+ naïve, memory Treg and T effector cells of RA patients (Walter et al., 2016). TGFβ PAS was higher in naïve Tregs of RA patients (p=0.016), while activity of the JAK-STAT3 pathway tended to increase in memory T effector cells of patients with RA (n=6, p=0.065, Figure 6), suggesting that these cell types contributed to the STP analysis results in whole blood samples, and may play a role in the pathogenesis of the disease.

**Figure 6.**
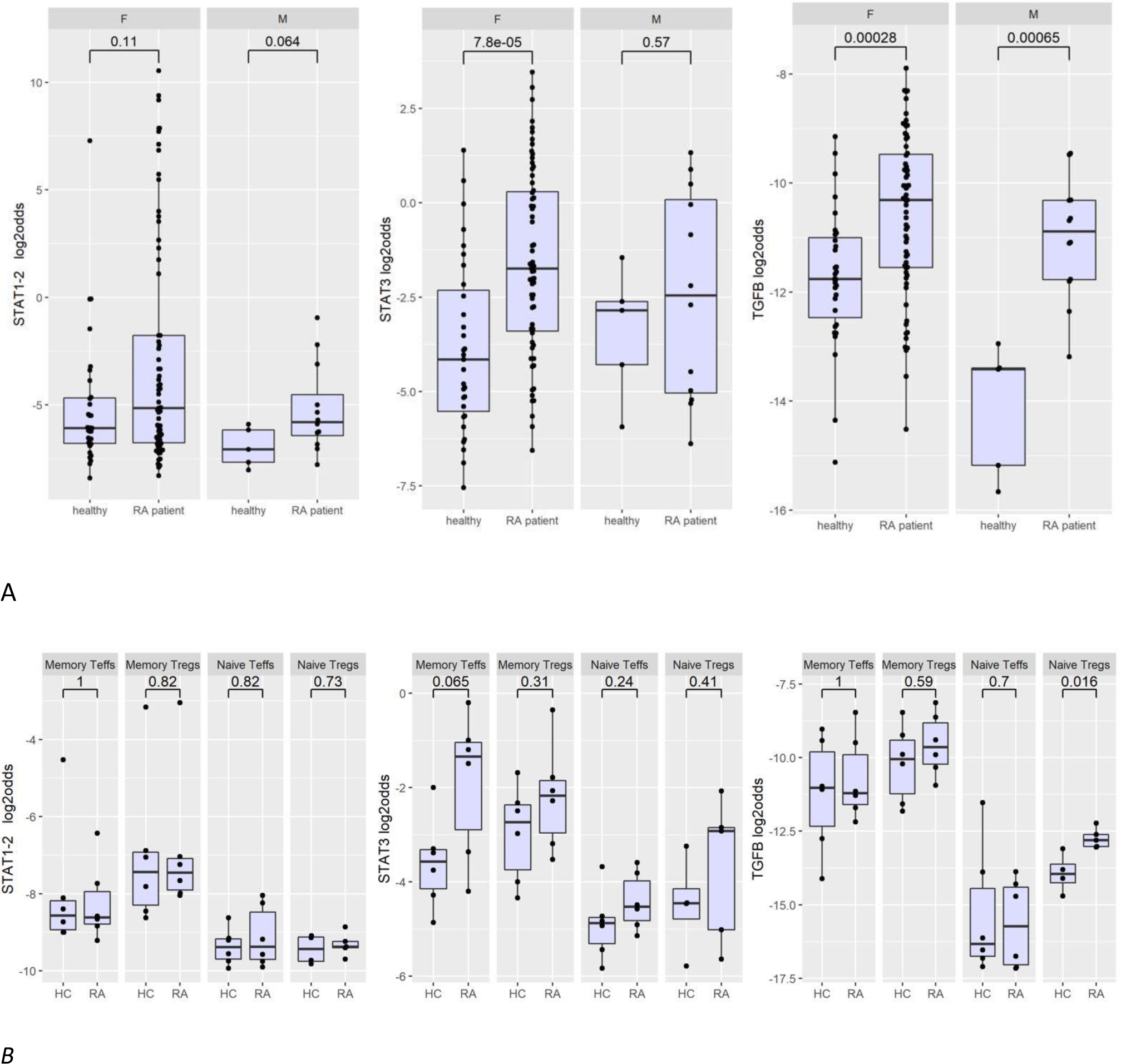
Clinical studies, patients with rheumatoid arthritis (RA) A. GSE93272 (Tasaki et al., 2018), whole blood samples from healthy volunteers and RA patients without prior treatment. JAK-STAT1/2, JAK-STAT3 and TGFβ PAS in log2 odds. F: female, M: Male B. GSE65010(Walter et al., 2016), CD4+ naïve and memory Treg and T effector cells of RA patients. HC: healthy control, RA: rheumatoid arthritis and Teffs: T effector cells. Boxplots were made using R. Exact p-values are indicated on top of the boxplots. The boxplot shows minimum, maximum, median, first quartile, third quartile and outliers which are plotted as individual points outside the minimum and maximum.

### Comparison of STP analysis with conventional Affymetrix data analysis

We compared our STP analysis results with the data analysis results reported in the publications associated with the public datasets (Supplementary information). Publicly available tools had been used to analyze the data, and comparison between sample groups revealed group-associated gene profiles and suggestions for differential involvement of cellular processes (not the same as signal transduction pathways). In contrast to the here presented STP analysis on single sample data, no results on analysis of individual patient samples were provided.

## Discussion

STP activity profiling of the different immune cell types revealed characteristic I-PAPs for each cell type in a resting and immune-activated state. Each I-PAP consists of combined quantitative pathway activity scores of the ER, AR, PI3K, MAPK, NFκB, JAK-STAT1/2, JAK-STAT3, TGFβ and Notch pathways. For some immune cell types more than one independent dataset was available for analysis, with largely similar results, providing support for reproducibility of the I-PAPs.

This is the first time that immune cells have been characterized by means of activity of nine major STPs that determine their function. All used STP assays have been validated before on multiple cell types for which pathway activity was known, supporting their use on different cell types including blood cells (Verhaegh et al., 2014), (van Ooijen et al., 2018), (Stolpe et al., 2019), (Canté-Barrett et al., 2020), (Bouwman et al., 2020), (Wesseling-Rozendaal et al., 2020). The remarkable similarity in pathway activities found between samples of healthy donors suggests strict regulation of signaling pathway activity and associated functional states of immune cells, in line with their important function in orchestrating immune responses.

Myeloid and lymphoid-derived cell types differed in their pathway activity profile, reflecting functional differences. While the PI3K growth factor pathway was more active in lymphocyte cell types, in myeloid derived cell types the MAPK growth factor pathway was most active, in combination with NFκB, TGFβ, and JAK-STAT3 pathways, in line with lineage-specific immune functions (Cantrell, 2015), (Newton and Dixit, 2012), (Shuai and Liu, 2003). The multiple pathway activity in myeloid cells is likely to reflect crosstalk between these signaling pathways (Chen and Ten Dijke, 2016),(Luo, 2017),(Shuai and Liu, 2003).

For a number of STPs specific functions in immune cells have been described, although generally not based on actual measurement of pathway activity but based on the expression levels of either ligands, receptors, or transcription factors associated with the pathway (e.g., see (Janghorban et al., 2018), (Batlle and Massagué, 2019). Our observation that PI3K activity in CD4+ and CD8+ T cells increased upon activation is in line with the role of this pathway to induce clonal proliferation of activated T cells, (Han et al., 2012). The observed inverse relationship of TGFβ pathway activity with CD4+ and CD8+ T cell activation supports our earlier findings that this pathway becomes active in CD4+ T cells upon exposure to breast cancer supernatant, inducing immunotolerance (Wesseling-Rozendaal et al., 2020). The immunosuppressive role of the TGFβ pathway in diseases such as cancer is well known (Li et al., 2007), (Yang, 2015),(Batlle and Massagué, 2019). Also, TGFβ pathway activity in neutrophils and macrophages is held responsible for inflammatory and tumor promoting functions (Futosi et al., 2013),(Fridlender et al., 2009), (Allavena et al., 2008).

The strong activation of the NFκB pathway induced by LPS and TNFα in myeloid cells, highlights the prominent role of this pathway as mediator of inflammatory immune responses by monocytes and neutrophils (Newton and Dixit, 2012),(Zhang et al., 2017). On the other hand, changes in NFκB pathway activity in lymphoid cells are more likely associated with other immune functions of this pathway, such as regulation of cell division, migration, antigen processing, and apoptosis, dependent on cellular context (Newton and Dixit, 2012), (Zhang et al., 2017).

The MAPK and Notch pathways have multiple context-dependent functions in the innate and adaptive immune response, and are involved in immune cell differentiation (Dong et al., 2002),(Janghorban et al., 2018). MAPK pathway activity was higher in Th1 than Th2 cells, in agreement with its involvement in Th1 differentiation (Dong et al., 2002). The high Notch pathway activity measured in B cells compared to T cells reflects its T cell-suppressing role in differentiation (Janghorban et al., 2018). Highest Notch pathway activity was associated with LPS and interferon-mediated activation of myeloid cell types, supportive of the role for this pathway in inflammatory responses (Gamrekelashvili et al., 2020).

We found the JAK-STAT pathways to be involved in many functional immune cell changes, in agreement with their distinct roles in a variety of immune reactions, dependent on cellular context. JAK-STAT1/2 pathway activity was generally induced by the presence of interferons, which are the typical ligands to activate this pathway (Shuai and Liu, 2003), (Barber, 2015). Indeed, we reported before increased JAK-STAT1/2 pathway activity in peripheral blood mononuclear cell (PBMC) samples during viral infections, known to be associated with strong interferon production (Bouwman et al., 2020). In the same study, JAK-STAT3 pathway activity increased selectively in PBMCs of patients with clinically severe infection and also after Bacillus Calmette-Guérin (BCG) boost vaccination (Bouwman et al., 2020). The current finding that JAK- STAT3 pathway activity was highest in activated cells of the monocytic lineage suggests that these cells were responsible for the earlier finding. We hypothesize that JAK-STAT3 pathway activity in monocytes (and macrophages) may reflect “trained immunity” which is associated with a strong inflammatory component; enabling direct measurement of this immune mechanism in patients (Netea et al., 2020), (Mulder et al., 2019).

AR pathway activity also increased in activated monocytes, in line with our earlier reporting on increased AR pathway activity in blood samples of patients with bacterial sepsis, in which monocytes play a dominant role (Bouwman et al., 2021), (Reyes et al., 2020). For the ER pathway low activity scores were consistently measured, with the only difference found between plasmacytoid and myeloid dendritic cells. Indeed, for both nuclear receptor (AR and ER) pathways, evidence is available for involvement in the innate immune response, with an immunosuppressive role for the AR pathway and a still controversial role for the ER pathway (Kovats, 2015), (Gubbels Bupp and Jorgensen, 2018), (Lai et al., 2012).

As referred to before, depending on cellular context individual STPs may crosstalk to involve other STPs to orchestrate the immune response. One way in which this may happen is illustrated by our time-related analysis of dendritic cell activation: viral infection sequentially activated the NFκB, JAK-STAT1/2, and finally the TGFβ pathway. The NFκB pathway is activated by the intracellular STING pathway to enable antigen processing, and subsequently induces production of interferons to activate the JAK-STAT1/2 pathway, necessary for presentation of antigens, while finally activating the TGFβ pathway, probably enabling DC migration (Barber, 2015),(Luo, 2017). *Analysis of samples from patients with rheumatoid arthritis*

In whole blood samples from patients with RA, activity of the TGFβ pathway and the JAK-STAT3 pathway were increased. Since whole blood consists of a mixture of cell types, these results are not directly informative on the functional activity state of the different immune cell types in the whole blood sample. Pathway analysis of a second clinical RA study (Treg and T effector cell samples) showed that at least the Treg cell population has abnormal STP activity: TGFβ pathway activity was higher in naïve Tregs, while JAK-STAT3 pathway tended to be more active in memory Tregs. Interesting, the investigators who generated the Affymetrix data of this study used Qlucore Omics Explorer software for the Affymetrix data analysis and concluded that no defect could be found in the Treg and T effector cells of the RA patients (Walter et al., 2016). However, both signaling pathways which we identified are known to play a pathogenic role in rheumatoid arthritis, and components of the JAK-STAT3 pathway (including its activating ligand IL-6) are important RA drug targets (Bottini et al., 2019), (Scott et al., 2010). The pro-inflammatory cytokine IL-6 and the immunosuppressive TGFβ together control differentiation between proinflammatory Th17 and immunosuppressive Treg cells, where IL-6 directs the balance towards Th17 cells by activating the JAK-STAT3 pathway, while TGFβ switches the balance towards Treg cells by activating the TGFβ pathway (Kimura and Kishimoto, 2010). Although insufficient to reveal the exact mechanism, we hypothesize that our findings reflect a disturbance in this balance, in favor of an inflammatory state. The variation in STP activity found in the RA patients suggests that the pathogenic mechanism varies among individual RA patients, suggesting that a personalized treatment approach is needed to provide the most effective treatment. Clinical use of our pathway assays may lie in diagnosis and monitoring of patients with RA, and in therapy response prediction: for example patients with high JAK-STAT3 pathway activity may be more likely to respond to JAK-STAT3 pathway inhibitors than patients with a JAK-STAT3 pathway activity in the normal range.

### Comparison of STP data analysis with conventionally used Affymetrix data analysis tools

We have shown in the current study that we can extract relevant biological information from transcriptome data that was not discovered by conventional data analysis, similar to our previous reports (Verhaegh et al., 2014), (van de Stolpe et al., 2021),(Bouwman et al., 2020), (Wesseling-Rozendaal et al., 2020),(Bouwman et al., 2021). Data analysis tools, such as GSEA and Ingenuity Pathway Analysis, provide differential involvement of all kinds of cellular processes when comparing two groups of samples and are suited for biomarker discovery. Our STP analysis is specifically designed to quantitatively measure STP activities (for a defined set of STPs) on an individual sample, for example for diagnostic purposes or to gain insight in immune cell biology. *STP analysis in perspective* We expect that measuring STP activity will enable functional quantification of the innate and adaptive immune response on blood samples of individual patients, or of a selected immune cell type. There are many clinically relevant diagnostic applications that can be developed, such as:

1. prediction and monitoring of response to cancer immunotherapy (see our publication (Wesseling-Rozendaal et al., 2020)), (2) differential diagnosis of viral versus bacterial infection, (see our publication (Bouwman et al., 2020)), (3) assessment and monitoring of the severity of a viral infection (Bouwman et al., 2020), (4) prediction of risk at development of sepsis in a patient with a bacterial infection, sepsis diagnosis and prognosis and prediction of treatment response (see our publication (Bouwman et al., 2021)), (5) diagnosis and treatment response prediction for many immune-mediated diseases (unpublished results, e.g. for Crohn’s Disease and Ulcerative Colitis, sarcoidosis, glomerulonephritis, psoriasis, pulmonary fibrosis); (6) prediction of late complications of kidney transplantation, such as fibrosis and glomerulonephritis (unpublished results). Determining and quantifying efficacy and side effects of vaccinations (see publication (Bouwman et al., 2020)) is another interesting application. Finally it is expected to improve development of immunomodulatory drugs, for multiple reasons: (1) The technology enables easy quantification of the effect of a drug/compound on STPs that determine immune cell functions;
2. Quantitative comparison between *in vitro* disease models and the actual disease in the patient with respect to the STP activity profile, enables choice of disease models that best represent *in vivo* disease; (3) Many existing drugs (for example developed for cancer applications) target these STPs, making them prime candidates for repurposing as immunomodulatory drugs.

To enable future implementation, it is necessary to have STP assays with shorter time-to-result than the Affymetrix microarray; for this reason the assay platform has been converted to qPCR, enabling a result within a few hours (www.philips.com/oncosignal). Clinical implementation will also require establishment of normal ranges for STP PAS, which is likely to be possible given the similarity of I-PAPs for healthy individual samples, and earlier reported results (Bouwman et al., 2020), (Bouwman et al., 2021). Another challenge is the type of blood sample to analyze. We have shown previously that STP analysis of mixed immune cell type samples, such as PBMCs or whole blood, may already provide clinically relevant results for a number of applications (Bouwman et al., 2021),(Bouwman et al., 2020). However, detailed analysis of the immune response of an individual will require isolation of specific immune cells of interest. While a single measurement can be directly informative, repeated analysis may add clinically relevant information by quantitative monitoring of changes in immune response.

## Conclusion

We have developed a new STP assay technology for quantitative measurement of the functional activity state of myeloid and lymphoid cell types, enabling for the first time characterization of the immune response in a patient sample and *in vitro*.

## Author contributions

Wilbert Bouwman: Data analysis, figures, writing

Wim Verhaegh: Pathway model development, data analysis, figures Arie van Doorn: Data analysis

Anja van de Stolpe: concept, data analysis, writing

Philips Molecular Pathway Dx (www.philips.com/oncosignal), Eindhoven, The Netherlands +31 6 12784841

