## Supplemental information for "Measuring host immune response status by simultaneous quantitative measurement of activity of signal transduction pathways that coordinate functional activity of immune cells from innate and adaptive immune system"

**Contains:**

I. STP activity analysis results, additional datasets for primary immune cell types
II. Comparison of Affymetrix dataset analysis results between STP analysis (this publication) and analysis performed as described by the investigators who originally generated the dataset

### I. Supplementary figures (STP activity analysis results, additional datasets for primary immune cell types)

| Group | Annotation per sample | AR | ER | FOXO | AP1 | NFKB | STAT1-2 | STAT3 | TGFB | NOTCH |
|---|---|---|---|---|---|---|---|---|---|---|
| 1. Monocytes | pool 1 | -13.2 | -6.9 | 11.1 | -4.9 | -4.5 | -6.6 | -6.9 | -9.4 | -3.1 |
| | pool 2 | -14.7 | -7.1 | 13.7 | -4.6 | 6.5 | -7.3 | -6.1 | -7.4 | -0.1 |
| | pool 3 | -14.0 | -6.2 | 13.7 | -4.2 | 10.4 | -6.4 | -4.5 | -8.2 | 2.2 |
| | pool 4 | -13.8 | -7.1 | 14.2 | -5.5 | 5.8 | -7.1 | -5.7 | -8.6 | 0.2 |
| | pool 5 | -12.7 | -5.2 | 16.6 | -4.9 | 8.1 | -7.4 | -5.3 | -8.5 | -0.7 |
| | pool 6 | -12.7 | -6.0 | 17.0 | -5.0 | 7.2 | -8.4 | -4.9 | -9.0 | -0.2 |
| | pool 7 | -12.3 | -6.9 | 10.6 | -6.4 | 0.6 | -6.5 | -7.1 | -12.1 | -2.5 |
| | pool 8 | -12.6 | -8.9 | 16.9 | -5.3 | -2.1 | -7.1 | -7.4 | -11.1 | -4.7 |
| | pool 9 | -11.6 | -9.5 | 18.2 | -5.5 | -4.2 | -7.0 | -7.0 | -10.2 | -4.0 |
| | pool 10 | -11.6 | -6.6 | 18.5 | -4.5 | -4.7 | -7.2 | -7.0 | -9.6 | -1.8 |
| 2. B- cells | pool 1 | -17.4 | -16.0 | 0.6 | -9.9 | -12.5 | -4.4 | -8.2 | -18.5 | -3.9 |
| | pool 2 | -18.1 | -16.7 | 0.5 | -9.2 | -14.6 | -4.8 | -11.7 | -18.0 | -4.4 |
| | pool 3 | -17.4 | -16.1 | -0.1 | -8.6 | -13.8 | -3.1 | -12.3 | -18.3 | -6.0 |
| | pool 4 | -17.8 | -16.1 | -0.1 | -9.5 | -13.9 | -4.3 | -10.8 | -19.4 | -7.1 |
| | pool 5 | -16.8 | -15.8 | 2.4 | -11.4 | -12.2 | -3.4 | -8.4 | -17.9 | -2.9 |
| 3. CD4+ T cells | pool 1 | -12.5 | -16.5 | -4.7 | -13.0 | -6.2 | -7.1 | -8.6 | -18.9 | -7.5 |
| | pool 2 | -11.2 | -14.3 | 10.9 | -8.4 | 1.7 | -8.4 | -3.8 | -11.8 | -3.1 |
| | pool 3 | -13.1 | -16.6 | -0.6 | -13.3 | -4.7 | -7.4 | -6.5 | -17.4 | -6.6 |
| | pool 4 | -12.0 | -16.1 | 2.7 | -15.0 | -5.5 | -7.8 | -5.7 | -18.0 | -6.3 |
| | pool 5 | -12.1 | -17.0 | -0.7 | -13.2 | -8.1 | -7.2 | -6.6 | -19.2 | -6.9 |
| 4. CD8+ T cells | pool 1 | -11.5 | -16.8 | 0.9 | -11.2 | -4.1 | -8.4 | -6.5 | -13.1 | -4.7 |
| | pool 2 | -12.2 | -16.8 | 0.3 | -11.7 | -3.7 | -8.1 | -5.2 | -14.0 | -5.9 |
| | pool 3 | -11.5 | -16.8 | -2.0 | -10.5 | -4.3 | -8.5 | -7.7 | -14.2 | -6.2 |
| | pool 4 | -16.1 | -17.1 | -6.9 | -12.8 | -6.2 | -7.1 | -9.7 | -18.3 | -5.7 |
| | pool 5 | -14.9 | -17.6 | -4.3 | -13.8 | -5.6 | -7.9 | -7.9 | -18.3 | -5.5 |
| 5. NK cells | pool 1 | -13.0 | -14.4 | -1.8 | -9.0 | -3.9 | -7.7 | -5.7 | -7.8 | -4.4 |
| | pool 2 | -15.0 | -15.8 | -5.9 | -11.5 | -1.4 | -7.0 | -8.4 | -11.4 | -5.3 |
| | pool 3 | -17.3 | -16.2 | -8.6 | -12.5 | -0.6 | -6.5 | -11.1 | -15.2 | -4.2 |
| | pool 4 | -15.5 | -15.6 | -6.5 | -12.6 | -1.4 | -7.1 | -8.3 | -10.9 | -5.5 |
| | pool 5 | -15.5 | -15.6 | -4.1 | -10.8 | -2.6 | -3.8 | -6.3 | -7.1 | -2.9 |
| 6. mDC | pool 1 | -13.3 | -10.9 | -4.9 | -6.2 | 5.6 | -5.7 | -7.4 | -15.9 | -7.3 |
| | pool 2 | -13.1 | -9.6 | -2.7 | -7.3 | 0.3 | -5.7 | -7.7 | -18.5 | -6.3 |
| | pool 3 | -14.3 | -11.8 | -9.0 | -12.5 | 0.8 | -5.0 | -7.5 | -17.5 | -8.0 |
| | pool 4 | -13.1 | -8.7 | -4.9 | -7.8 | 3.6 | -6.0 | -5.1 | -17.8 | -6.2 |
| | pool 5 | -13.5 | -11.3 | -4.5 | -8.4 | -0.8 | -6.8 | -6.5 | -18.7 | -7.2 |
| 7. pDC | pool 1 | -8.6 | -14.8 | -4.3 | -9.6 | -11.7 | 0.5 | -8.7 | -18.1 | -9.0 |
| | pool 2 | -8.3 | -14.9 | -3.6 | -10.2 | -12.4 | 1.8 | -8.5 | -19.4 | -8.1 |
| | pool 3 | -8.0 | -13.3 | -5.8 | -12.1 | -7.7 | 0.5 | -9.7 | -19.2 | -8.3 |
| | pool 4 | -7.4 | -14.0 | -2.3 | -12.5 | -14.7 | 0.0 | -8.7 | -18.7 | -8.6 |
| | pool 5 | -7.5 | -13.4 | -2.6 | -12.2 | -13.1 | -0.9 | -9.2 | -19.0 | -8.1 |

| | | | | | | | | | | |
|---|---|---|---|---|---|---|---|---|---|---|
| 1,6,7 vs. 2-5 | delta mean | -2.7 | -6.5 | -6.7 | -3.9 | -4.8 | -1.6 | -0.9 | -1.6 | -0.7 |
| | t-test p-value | 0.001085 | 1.13E-08 | 0.012667 | 2.33E-05 | 0.022999 | 0.065072 | 0.137261 | 0.262859 | 0.448545 |
| 1 vs. 6,7 | delta mean | 2.2 | -5.2 | -19.5 | -4.8 | -7.3 | 4.4 | -1.7 | -8.9 | -6.2 |
| | t-test p-value | 0.046827 | 9.11E-06 | 4.9E-12 | 9.41E-05 | 0.028843 | 0.002592 | 0.005826 | 1.84E-11 | 1.58E-06 |
| 2 vs. 3,4,5 | delta mean | 3.9 | -0.1 | -2.7 | -2.2 | 9.6 | -3.3 | 3.1 | 4.1 | -0.5 |
| | t-test p-value | 1.35E-06 | 0.776482 | 0.054636 | 0.005041 | 8.33E-10 | 6.92E-06 | 0.016062 | 0.001554 | 0.55408 |
| 3 vs. 4 | delta mean | -1.1 | -0.9 | -3.9 | 0.6 | -0.2 | -0.4 | -1.2 | 1.5 | 0.5 |
| | t-test p-value | 0.337958 | 0.125923 | 0.240733 | 0.645094 | 0.906057 | 0.267641 | 0.316722 | 0.419832 | 0.579261 |
| 5 vs. 2,3,4 | delta mean | 1.0 | -0.9 | 5.3 | -0.2 | -5.6 | -0.1 | 0.0 | -6.5 | -1.1 |
| | t-test p-value | 0.341245 | 0.034323 | 0.005395 | 0.859121 | 0.000664 | 0.903656 | 0.992236 | 0.00715 | 0.099683 |
| 2 vs. 3,4,5 | delta mean | 3.9 | -0.1 | -2.7 | -2.2 | 9.6 | -3.3 | 3.1 | 4.1 | -0.5 |
| | t-test p-value | 1.35E-06 | 0.776482 | 0.054636 | 0.005041 | 8.33E-10 | 6.92E-06 | 0.016062 | 0.001554 | 0.55408 |
| 6 vs. 7 | delta mean | 5.5 | -3.6 | 1.5 | -2.9 | -13.8 | 6.2 | -2.1 | -1.2 | -1.4 |
| | t-test p-value | 1.29E-07 | 0.001052 | 0.259737 | 0.055395 | 3.1E-05 | 7.09E-06 | 0.008227 | 0.07156 | 0.00948 |

***S1.*** *GSE28490, (Allantaz et al., 2012). Immune blood cell types (neutrophils, eosinophils, monocytes, B cells, NK cells, CD4 T cells, CD8 T cells, mDCs and pDCs) isolated from peripheral blood from healthy volunteers. Pathway activity scores are depicted in log2odds values. Color coding visualizes pathway activity scores ranging from blue (lowest) to red (highest pathway activity scores). P<0.01, green; p<0.5 orange.*

| Group | AR | ER | FOXO | AP1 | NFKB | STAT1-2 | STAT3 | TGFB | NOTCH |
|---|---|---|---|---|---|---|---|---|---|
| Resting NK | -7.4 | -11.0 | 7.8 | -9.5 | 8.5 | -6.8 | 0.2 | -2.9 | 0.3 |
| NK 2 hrs IL-2 activation | -8.4 | -11.4 | 1.5 | -6.8 | 2.5 | -6.9 | -3.0 | -1.0 | -1.6 |
| NK 8 hrs IL-2 activation | -16.1 | -10.1 | -6.8 | -8.9 | -0.4 | -7.1 | -5.1 | -8.4 | -4.5 |
| NK 24 hrs IL-2 activation | -13.3 | -9.6 | -4.9 | -11.3 | 7.6 | -7.4 | 3.8 | -7.1 | -5.1 |

A

| Group | AR | ER | FOXO | AP1 | NFKB | STAT1-2 | STAT3 | TGFB | NOTCH |
|---|---|---|---|---|---|---|---|---|---|
| stimulated_24hr_donor_1 | -14.0 | -13.9 | 8.0 | -14.0 | 8.2 | -4.2 | 10.3 | -7.6 | -4.5 |
| stimulated_24hr_donor_2 | -13.8 | -13.3 | 7.3 | -14.1 | 9.7 | -6.0 | 12.6 | -8.3 | -4.3 |
| unstimulated_freshly_isolated_donor_1 | -14.4 | -16.4 | 4.1 | -14.2 | 0.6 | -6.7 | -7.4 | -9.6 | -3.2 |
| unstimulated_freshly_isolated_donor_2 | -8.9 | -15.6 | 4.7 | -14.4 | 0.3 | -6.3 | -7.3 | -10.3 | -3.6 |

B

***S2.*** *Natural killer cells,*

*A) GSE8059, (Dybkaer et al., 2007). NK-cells resting and activated with IL-2 (100 IU/ml) for 2, 8 and 24h.*

*B) GSE22919,(Smith et al., 2010). NK cells unstimulated and stimulated with IL-2 (100U/mL) + IL-12 (10ng/mL) and IL-18 (100ng/mL).*

| Group | Annotation per sample | AR | ER | FOXO | AP1 | NFKB | STAT1-2 | STAT3 | TGFB | NOTCH |
|---|---|---|---|---|---|---|---|---|---|---|
| 1. Saline | subject 1 | -6.1 | -11.8 | 0.1 | -3.2 | 6.7 | -4.3 | 4.2 | -7.1 | -3.3 |
| | subject 2 | -5.3 | -11.9 | -0.5 | -4.2 | -3.1 | -4.5 | -1.3 | -8.9 | -4.5 |
| | subject 3 | -6.3 | -10.5 | 5.7 | -5.0 | 6.1 | -4.2 | 3.3 | -6.7 | -4.1 |
| | subject 5 | -6.1 | -11.1 | 4.2 | -4.4 | 11.8 | -5.2 | 3.5 | -4.2 | -1.0 |
| | subject 6 | -6.1 | -10.1 | 4.7 | -5.0 | 6.1 | -6.3 | -0.6 | -6.3 | -4.9 |
| | subject 7 | -6.5 | -11.8 | 2.2 | -4.1 | 9.3 | -5.2 | 3.7 | -5.7 | -1.6 |
| 2. LPS | subject 1 | -5.3 | -12.5 | 1.2 | -3.0 | 18.9 | -2.6 | 5.6 | -4.2 | -2.4 |
| | subject 3 | -6.4 | -12.1 | 1.7 | -3.5 | 23.1 | 2.3 | 9.2 | -5.1 | -2.5 |
| | subject 4 | -6.2 | -12.0 | 1.1 | -4.3 | 21.0 | -2.1 | 6.1 | -4.8 | -2.0 |
| | subject 5 | -5.8 | -12.0 | 1.0 | -0.3 | 22.7 | -1.1 | 9.5 | -1.4 | 0.5 |
| | subject 6 | -6.2 | -12.0 | 0.7 | -2.3 | 22.3 | -0.8 | 8.7 | -4.1 | -0.5 |
| | subject 7 | -6.1 | -13.4 | 0.8 | -1.7 | 23.1 | -0.5 | 9.9 | -3.8 | -0.7 |

| | | | | | | | | | | |
|---|---|---|---|---|---|---|---|---|---|---|
| 1 vs. 2 | delta mean | 0.0 | 1.4 | 2.0 | -1.5 | -15.1 | -4.2 | -5.4 | -2.3 | -1.8 |
| | t-test p-value | 0.985001 | 0.002802 | 0.068968 | 0.031379 | 3.56E-05 | 0.000156 | 0.000493 | 0.008023 | 0.029465 |

***S3.*** *GSE40885, (Reynier et al., 2012). Alveolar macrophages obtained from a contralateral lung segment challenged with LPS (4 ng/kg body weight) or saline (control).*

| Group | Annotation per sample | AR | ER | FOXO | AP1 | NFKB | STAT1-2 | STAT3 | TGFB | NOTCH |
|---|---|---|---|---|---|---|---|---|---|---|
| 1. CD4+ naïve | Replicate 2 | -11.5 | -20.4 | 5.2 | -17.7 | 4.0 | -8.1 | -8.0 | -13.1 | -2.3 |
| | Replicate 3 | -11.8 | -20.3 | 0.4 | -18.1 | -1.5 | -6.9 | -8.5 | -13.0 | -2.8 |
| 2. CD4+ activated | Replicate 1 | -9.4 | -19.8 | -7.0 | -18.0 | 0.1 | -1.6 | -5.1 | -16.1 | -2.6 |
| | Replicate 2 | -9.5 | -19.8 | -6.8 | -16.9 | -0.1 | -1.7 | -6.6 | -16.4 | -2.1 |
| | Replicate 3 | -12.6 | -19.6 | -6.1 | -15.4 | -1.6 | -1.5 | -9.3 | -14.8 | -0.9 |
| 3. CD4+ activated + IL12 (Th1) | Replicate 1 | -8.3 | -19.6 | -5.3 | -19.4 | -0.1 | -2.7 | -2.5 | -15.5 | -3.6 |
| | Replicate 2 | -7.4 | -19.5 | -3.3 | -18.4 | -0.5 | -2.1 | -2.4 | -15.8 | -2.4 |
| | Replicate 3 | -8.8 | -19.4 | -3.7 | -15.9 | -1.7 | -3.3 | -6.6 | -13.1 | -2.8 |
| 4. CD4+ activated + IL4 (Th2) | Replicate 1 | -10.4 | -19.5 | -3.1 | -16.8 | -7.1 | -6.0 | -6.7 | -19.7 | -3.0 |
| | Replicate 2 | -11.0 | -20.2 | -5.9 | -15.7 | -2.0 | -4.5 | -8.4 | -18.8 | -2.9 |
| | Replicate 3 | -12.7 | -19.5 | 1.0 | -16.1 | -3.0 | -6.0 | -9.6 | -16.2 | -2.2 |

| | | | | | | | | | | |
|---|---|---|---|---|---|---|---|---|---|---|
| 1 vs. 2 | delta mean | 1.2 | 0.6 | -9.5 | 1.1 | -1.8 | 5.9 | 1.3 | -2.7 | 0.7 |
| | t-test p-value | 0.387111 | 0.009662 | 0.155687 | 0.266907 | 0.631143 | 0.066395 | 0.410624 | 0.030814 | 0.295629 |
| 2 vs. 3,4 | delta mean | 0.7 | 0.1 | 3.2 | -0.3 | -1.9 | -2.5 | 1.0 | -0.7 | -0.9 |
| | t-test p-value | 0.617402 | 0.354089 | 0.021409 | 0.773915 | 0.157354 | 0.014281 | 0.601685 | 0.519215 | 0.18578 |
| 2 vs. 3 | delta mean | 2.3 | 0.3 | 2.5 | -1.1 | -0.2 | -1.1 | 3.2 | 1.0 | -1.0 |
| | t-test p-value | 0.149679 | 0.032405 | 0.040751 | 0.431792 | 0.768251 | 0.085709 | 0.168605 | 0.390881 | 0.167513 |
| 2 vs. 4 | delta mean | -0.9 | 0.0 | 4.0 | 0.5 | -3.5 | -3.9 | -1.2 | -2.5 | -0.8 |
| | t-test p-value | 0.525597 | 0.96902 | 0.184881 | 0.553179 | 0.146009 | 0.016446 | 0.470216 | 0.12637 | 0.227599 |
| 1 vs. 3,4 | delta mean | 1.9 | 0.7 | -6.2 | 0.8 | -3.6 | 3.4 | 2.3 | -3.5 | -0.2 |
| | t-test p-value | 0.064911 | 0.001644 | 0.197569 | 0.243846 | 0.392127 | 0.024193 | 0.12428 | 0.016961 | 0.528564 |
| 1 vs. 3 | delta mean | 3.5 | 0.9 | -6.9 | 0.0 | -2.0 | 4.8 | 4.4 | -1.7 | -0.3 |
| | t-test p-value | 0.008654 | 0.002995 | 0.195501 | 0.99647 | 0.595507 | 0.03472 | 0.081299 | 0.182828 | 0.493579 |
| 1 vs. 4 | delta mean | 0.3 | 0.6 | -5.5 | 1.7 | -5.3 | 2.0 | 0.1 | -5.2 | -0.1 |
| | t-test p-value | 0.717226 | 0.118896 | 0.206303 | 0.020263 | 0.259857 | 0.115904 | 0.93643 | 0.038183 | 0.739914 |
| 3 vs. 4 | delta mean | -3.2 | -0.2 | 1.4 | 1.7 | -3.3 | -2.8 | -4.4 | -3.5 | 0.2 |
| | t-test p-value | 0.02134 | 0.401222 | 0.552599 | 0.244772 | 0.165828 | 0.014521 | 0.067632 | 0.065249 | 0.674315 |

***S4.*** *GSE32959, (Aijö et al., 2012). Naïve CD4+ cells, CD4+ activated (antiCD3 500 ng/well+ antiCD28 (500 ng/ml), CD4+ activated + Th1 polarization (IL-12 (2.5ng/ml)) and CD4+ activated + Th2 polarization (IL-4 (10ng/ml and anti-IL-12 (10µg/ml). For induction of Th1 cell polarization, IL-12 (2.5 ng/ml) was added to the cultures. At 48h after activation, IL-2 was added (17 ng/ml) to all the cells and the polarizing conditions were maintained throughout the culture.*

| Group | AR | ER | FOXO | AP1 | NFKB | STAT1-2 | STAT3 | TGFB | NOTCH |
|---|---|---|---|---|---|---|---|---|---|
| Primary peripheral blood B cell - Resting | -8.95 | -14.50 | 8.57 | -10.31 | 0.24 | -4.53 | -2.65 | -6.73 | 2.43 |
| Primary peripheral blood B cell - Resting | -9.19 | -15.22 | 8.42 | -10.51 | 0.54 | -4.81 | -3.76 | -6.61 | 1.38 |
| Primary peripheral blood B cell + antiIgM -1hr | -6.20 | -12.73 | 6.35 | -10.14 | 5.82 | -2.53 | 5.58 | -2.92 | 4.62 |
| Primary peripheral blood B cell + antiIgM -3hr | -6.90 | -13.38 | 1.70 | -9.17 | 3.30 | -6.92 | 2.59 | -6.68 | 5.08 |

***S5.*** *GSE9119, (Shaffer et al., 2008). B-cells resting and activated with anti IgM (25ug/ml) for 1hr and 3hrs.*

*Note: 3 samples failed our Affymetrix QC, on Cmoff criterium.*

| infection | Donor | Time (h) | AR | ER | FOXO | AP1 | NFKB | STAT1-2 | STAT3 | TGFB | NOTCH |
|---|---|---|---|---|---|---|---|---|---|---|---|
| AF | Donor 1, replicate 1 | 0 | -7.5 | -9.4 | 1.1 | -8.9 | 12.8 | -11.5 | 0.7 | -15.4 | -3.3 |
| AF | Donor 1, replicate 1 | 0 | -9.5 | -11.3 | -0.9 | -9.5 | 12.1 | -11.3 | -4.0 | -14.4 | -3.6 |
| AF | Donor 1, replicate 2 | 0 | -12.4 | -13.4 | -1.3 | -7.1 | 14.7 | -11.0 | -3.6 | -15.9 | -5.1 |
| AF | Donor 2, replicate 2 | 0 | -9.3 | -10.4 | -2.5 | -8.8 | 9.6 | -11.3 | -0.9 | -14.2 | -4.2 |
| AF | Donor 2, replicate 1 | 1 | -10.0 | -11.8 | -1.9 | -9.4 | 13.3 | -11.0 | -2.6 | -14.3 | -4.2 |
| AF | Donor 1, replicate 2 | 1 | -12.4 | -13.9 | -0.4 | -8.1 | 14.2 | -10.1 | -3.2 | -16.1 | -2.9 |
| NDV | Donor 1, replicate 1 | 1 | -7.3 | -10.1 | 3.3 | -8.0 | 13.6 | -11.4 | 1.6 | -14.2 | -3.2 |
| NDV | Donor 2, replicate 1 | 1 | -9.9 | -11.0 | -0.8 | -9.8 | 12.2 | -10.9 | -3.9 | -14.5 | -4.5 |
| NDV | Donor 1, replicate 2 | 1 | -11.6 | -13.5 | -0.3 | -8.5 | 15.8 | -10.5 | -3.0 | -16.0 | -2.4 |
| NDV | Donor 2, replicate 2 | 1 | -9.1 | -11.5 | 3.1 | -9.2 | 15.0 | -11.2 | -1.6 | -14.3 | -2.5 |
| AF | Donor 2, replicate 1 | 2 | -9.6 | -12.1 | -0.2 | -10.3 | 12.9 | -11.3 | -2.9 | -14.4 | -4.2 |
| AF | Donor 1, replicate 2 | 2 | -11.4 | -13.8 | 0.1 | -6.9 | 12.0 | -10.4 | -2.4 | -14.4 | -4.6 |
| NDV | Donor 1, replicate 1 | 2 | -7.2 | -9.3 | 3.0 | -8.5 | 16.7 | -10.5 | 2.4 | -10.1 | -3.6 |
| NDV | Donor 2, replicate 1 | 2 | -11.5 | -11.7 | -2.9 | -11.5 | 10.5 | -10.9 | -4.6 | -13.4 | -3.8 |
| NDV | Donor 1, replicate 2 | 2 | -11.5 | -13.5 | 0.2 | -8.5 | 16.3 | -10.5 | -2.2 | -14.0 | -4.7 |
| NDV | Donor 2, replicate 2 | 2 | -9.0 | -11.8 | 5.3 | -9.1 | 15.8 | -10.6 | -1.5 | -12.5 | -2.6 |
| NDV | Donor 1, replicate 1 | 4 | -6.8 | -11.1 | 8.9 | -9.1 | 20.1 | -0.9 | 2.4 | -12.0 | -3.5 |
| NDV | Donor 2, replicate 1 | 4 | -10.2 | -11.4 | -0.2 | -11.5 | 15.0 | -7.4 | -4.3 | -12.8 | -3.4 |
| NDV | Donor 1, replicate 2 | 4 | -12.4 | -13.9 | 2.4 | -8.6 | 19.6 | -4.7 | -2.0 | -13.3 | -2.9 |
| NDV | Donor 2, replicate 2 | 4 | -9.0 | -13.8 | 10.3 | -10.3 | 22.1 | -0.6 | -0.7 | -11.5 | -2.2 |
| AF | Donor 1, replicate 1 | 6 | -6.4 | -9.8 | 4.3 | -8.2 | 14.7 | -10.0 | 0.3 | -11.1 | -3.2 |
| AF | Donor 1, replicate 2 | 6 | -12.5 | -13.5 | 2.7 | -6.0 | 12.0 | -9.0 | -1.2 | -12.9 | -4.6 |
| AF | Donor 2, replicate 2 | 6 | -8.1 | -12.3 | 11.1 | -6.3 | 15.2 | -9.6 | -0.8 | -10.8 | -3.1 |
| NDV | Donor 1, replicate 1 | 6 | -6.6 | -10.9 | 8.6 | -9.8 | 21.8 | 2.5 | 2.5 | -11.7 | -1.4 |
| NDV | Donor 1, replicate 2 | 6 | -10.0 | -15.8 | 6.9 | -8.2 | 28.1 | 1.4 | -1.8 | -14.2 | -2.3 |
| NDV | Donor 2, replicate 2 | 6 | -7.9 | -13.9 | 12.1 | -10.0 | 27.3 | 2.0 | 0.9 | -12.0 | -2.7 |
| NDV | Donor 1, replicate 1 | 8 | -6.5 | -11.1 | 9.4 | -10.8 | 23.6 | 4.7 | 2.7 | -10.6 | -2.1 |
| NDV | Donor 2, replicate 1 | 8 | -11.2 | -13.0 | 5.1 | -12.7 | 23.5 | -0.4 | -2.3 | -13.4 | -4.0 |
| NDV | Donor 1, replicate 2 | 8 | -12.1 | -14.5 | 8.6 | -10.5 | 28.6 | 2.6 | -0.6 | -10.4 | -3.7 |
| NDV | Donor 2, replicate 2 | 8 | -7.0 | -15.4 | 11.1 | -7.8 | 29.9 | 4.5 | 1.4 | -5.8 | -2.5 |
| AF | Donor 1, replicate 1 | 10 | -6.5 | -10.2 | 6.7 | -6.6 | 17.1 | -10.4 | 1.7 | -9.3 | -3.3 |
| AF | Donor 2, replicate 1 | 10 | -10.2 | -12.1 | 0.2 | -11.0 | 12.4 | -10.1 | -3.8 | -12.8 | -3.8 |
| AF | Donor 1, replicate 2 | 10 | -9.7 | -11.3 | 5.0 | -5.9 | 14.5 | -9.7 | 0.4 | -9.9 | -3.8 |
| AF | Donor 2, replicate 2 | 10 | -7.7 | -12.5 | 10.2 | -7.2 | 14.3 | -9.5 | -0.4 | -8.6 | -3.2 |
| NDV | Donor 1, replicate 1 | 10 | -6.8 | -12.3 | 11.0 | -11.4 | 25.9 | 5.5 | 5.7 | -4.2 | -2.2 |
| NDV | Donor 1, replicate 2 | 10 | -9.7 | -13.5 | 10.9 | -9.0 | 31.1 | 5.5 | 1.0 | -4.6 | -4.0 |
| NDV | Donor 2, replicate 2 | 10 | -7.8 | -14.0 | 10.7 | -11.7 | 30.2 | 6.7 | 2.5 | -6.8 | -3.1 |
| NDV | Donor 1, replicate 1 | 12 | -6.9 | -10.7 | 13.6 | -11.0 | 23.8 | 7.4 | 6.7 | -2.2 | -1.4 |
| NDV | Donor 1, replicate 2 | 12 | -8.5 | -12.9 | 10.9 | -10.8 | 31.6 | 6.4 | 1.3 | -3.1 | -0.5 |
| NDV | Donor 2, replicate 2 | 12 | -7.1 | -11.2 | 9.6 | -8.7 | 32.1 | 9.1 | 3.2 | -3.3 | -2.5 |
| NDV | Donor 1, replicate 1 | 14 | -6.7 | -11.0 | 13.0 | -11.2 | 23.9 | 8.4 | 6.7 | -1.6 | -1.7 |
| NDV | Donor 2, replicate 1 | 14 | -11.1 | -14.5 | 2.4 | -11.5 | 21.5 | 0.8 | -2.6 | -8.7 | -0.9 |
| NDV | Donor 1, replicate 2 | 14 | -8.7 | -12.7 | 12.8 | -10.8 | 33.0 | 7.7 | 2.4 | -1.3 | -3.3 |
| NDV | Donor 2, replicate 2 | 14 | -8.1 | -11.9 | 9.7 | -8.3 | 32.0 | 9.3 | 2.9 | -3.2 | -1.6 |
| NDV | Donor 1, replicate 1 | 16 | -6.7 | -11.0 | 15.9 | -10.6 | 25.0 | 9.1 | 5.9 | -2.5 | -1.7 |
| NDV | Donor 2, replicate 1 | 16 | -11.3 | -14.9 | 6.1 | -12.5 | 21.9 | 2.2 | -3.1 | -8.1 | -1.5 |
| NDV | Donor 1, replicate 2 | 16 | -8.3 | -11.6 | 13.9 | -10.3 | 32.8 | 6.7 | 1.6 | -1.2 | -1.6 |
| NDV | Donor 2, replicate 2 | 16 | -9.1 | -15.0 | 7.0 | -9.4 | 28.2 | 6.7 | 3.3 | -6.2 | -2.3 |
| AF | Donor 2, replicate 1 | 18 | -10.0 | -12.6 | 0.7 | -12.2 | 10.6 | -10.0 | -4.4 | -11.4 | -3.6 |
| AF | Donor 1, replicate 2 | 18 | -8.7 | -12.2 | 9.5 | -6.2 | 17.9 | -9.2 | -0.6 | -11.7 | -4.3 |
| NDV | Donor 1, replicate 2 | 18 | -8.6 | -12.9 | 13.4 | -10.4 | 32.3 | 7.8 | 1.7 | -1.7 | -2.1 |
| NDV | Donor 2, replicate 2 | 18 | -8.7 | -14.3 | 11.0 | -11.2 | 27.9 | 8.2 | 3.4 | -4.9 | -2.2 |

A

| Group | Annotation per sample | AR | ER | FOXO | AP1 | NFKB | STAT1-2 | STAT3 | TGFB | NOTCH |
|---|---|---|---|---|---|---|---|---|---|---|
| 1. Immature DCs | replicate 1 | -7.6 | -6.6 | 5.2 | -9.4 | 14.2 | -10.6 | 6.0 | -12.3 | 0.7 |
| | replicate 2 | -7.6 | -6.2 | 6.2 | -8.9 | 14.2 | -10.6 | 7.5 | -10.9 | 0.5 |
| | replicate 3 | -7.4 | -6.1 | 6.0 | -7.3 | 13.7 | -10.5 | 7.2 | -11.6 | 0.5 |
| 2. Maturing DCs | replicate 1 | -7.4 | -9.5 | 16.4 | -0.7 | 27.6 | 6.3 | 10.2 | 0.9 | 3.0 |
| | replicate 2 | -7.2 | -9.0 | 18.0 | -0.7 | 24.9 | 6.9 | 10.7 | 2.4 | 3.4 |
| | replicate 3 | -6.9 | -9.3 | 17.3 | -0.8 | 26.5 | 7.6 | 10.7 | 0.5 | 2.8 |
| 3. Tolerogenic DCs | replicate 1 | -7.0 | -8.3 | 10.2 | -6.3 | 10.1 | -6.5 | 3.7 | -13.0 | 0.5 |
| | replicate 2 | -7.1 | -8.0 | 11.1 | -5.3 | 12.2 | -6.8 | 5.2 | -11.9 | 0.3 |
| | replicate 3 | -7.0 | -8.1 | 10.2 | -6.0 | 10.2 | -6.7 | 4.7 | -11.8 | 0.9 |
| 4. Activated tolerogenic DCs | replicate 1 | -6.9 | -9.2 | 14.6 | -2.0 | 28.1 | 3.4 | 12.1 | -2.8 | 3.6 |
| | replicate 2 | -6.5 | -9.0 | 15.9 | -1.1 | 28.2 | 3.2 | 11.8 | -0.9 | 5.0 |
| | replicate 3 | -6.4 | -8.5 | 16.6 | -1.0 | 28.1 | 3.0 | 10.9 | -1.3 | 4.4 |

| | | | | | | | | | | |
|---|---|---|---|---|---|---|---|---|---|---|
| 1 vs 2 | delta mean | -0.4 | 3.0 | -1.4 | -11.5 | -7.8 | -12.3 | -17.5 | -3.6 | -12.9 | -2.5 |
| | t-test p-value | 0.122031 | 0.000143 | 0.024902 | 7.41E-05 | 0.006225 | 0.003089 | 0.000413 | 0.009235 | 0.000106 | 0.002872 |
| 3 vs 4 | delta mean | -0.4 | 0.8 | -4.0 | -5.2 | -4.5 | -17.3 | -9.9 | -7.0 | -10.6 | -3.7 |
| | t-test p-value | 0.081511 | 0.051512 | 1.41E-05 | 0.004596 | 0.000497 | 0.001583 | 1.59E-06 | 0.000324 | 0.000246 | 0.004037 |

B

***S6.** Dendritic cells*

*A. GSE18791, (Zaslavsky et al., 2010a). All STP PAS from this study. Dendritic cells (Monocyte-derived DCs) from two donors, resting and activated by infection with Newcastle disease virus (NDV), at 1, 2, 4, 6, 8, 10, 12, 14, 16 and 18 hours after infection versus control.*

*B) GSE23371, (Jansen et al., 2011); DCs immature (IL4 and GM-CSF), mature (IL4 and GM-CSF + 6h LPS), tolerogenic (IL4 and GM-CSF + 24 hr IL10/dexamethasone), activated tolerogenic (IL4 and GM-CSF + IL10/dexamethasone 24 hr, then LPS 6 hr).*

### II. Comparison of Affymetrix dataset analysis results between STP analysis (this publication) and analysis performed as described by the investigators who originally generated the dataset

GSE72642, (Du et al., 2006)

Data analysis: Differential gene expression. No Pathway analysis

No signaling pathway activity identified.

GSE15743, (Stegmann et al., 2010)

Data analysis: Differential expression of TRAIL gene. No pathway analysis.

No signaling pathway activity identified.

GSE22103, (Kotz et al., 2010)

Data analysis: Ingenuity Pathway analysis.

Identified cellular processes ("pathways") (top 10):

- Upregulated
    - Oxidative phosphorylation
    - Mitochondrial dysfunction
    - Ubiquinone biosynthesis
    - Polyamine regulation in colon cancer
    - Protein ubiquitination pathway
    - Inositol metabolism
    - Pyrimidine metabolism
    - RAN signaling
    - Mitotic roles of polo-like kinases

- N-glycan biosynthesis

- Downregulated:
    - Antigen presentation pathway
    - IL-4 signaling
    - GNRH signaling
    - Retinoic acid-mediated apoptosis signaling
    - Interferon signaling
    - Corticotropin releasing hormone signaling
    - Death receptor signaling
    - B cell receptor signaling
    - Huntington's disease signaling
    - IL-3 signaling

GSE38351 (Smiljanovic et al., 2012)

Data analysis: Ingenuity Pathway Analysis

JAK-STAT1/2 pathway identified, no information on activity of the pathway.

GSE43596, (Lowe et al., 2014).

Data analysis: differential gene expression.

P53/NFκB transcription factor coregulation of immune response genes. No information on activity of the NFκB pathway.

GSE71566, (Kanduri et al., 2015)

Data analysis: differential gene expression. No Pathway analysis.

No signaling pathways identified.

GSE63129, (Maeda et al., 2014)

Data analysis: differential expression. No pathway analysis.

No signaling pathways identified.

GSE39411, (Vallat et al., 2013)

Data analysis: various methods(differential expression, regression, gene networks). No pathway analysis.

No signaling pathways identified for activation of B cells.

GSE18791, (Zaslavsky et al., 2010b)

Data analysis: TRANSFAC transcription factor heatmap construction.

Identification of transcriptional cascade: Interferon response factors (IRF), STATs and NFκB (STAT activity preceded NFκB in the TRANSFAC network).

No identification of signal transduction pathway activity.

GSE65010 (Walter et al., 2016).

Data analysis: Qlucore Omics Explorer software, version 3.0.

Result: no statistically significant differences between RA patients and healthy controls.

Conclusion, cited from abstract: *"Our findings indicate that there is no global defect in either CD45RO1 or CD45RA1 Treg cells in the PB of patients with chronic RA".*

GSE93272, (Tasaki et al., 2018)

Data analysis: Differential expression and ssGSEA for pathway analysis

Identified cellular processes (“pathways”):

- Regulation of complement cascade
- Coagulation
- COMP pathway
- Initial triggering of complement
- Complement cascade
- Telomerase pathway
- CXCR3 pathway

No signal transduction pathways identified.
